# Locally-correlated kinetics of post-replication DNA methylation reveals processivity and region-specificity in DNA methylation maintenance

**DOI:** 10.1101/2021.09.28.462223

**Authors:** Honglei Ren, Robert B. Taylor, Timothy L. Downing, Elizabeth L. Read

## Abstract

DNA methylation occurs predominantly on cytosine-phosphate-guanine (CpG) dinucleotides in the mammalian genome, and the methylation landscape is maintained over mitotic cell division. It has been posited that coupling of maintenance methylation activity among neighboring CpGs is critical to collective stability over cellular generations, however the mechanism of this coupling is unclear. We used mathematical models and stochastic simulation to analyze data from experiments that probe genome-wide methylation of nascent DNA postreplication in cells. We find that DNA methylation maintenance rates on individual CpGs are locally correlated, and the degree of this correlation varies by genomic regional context. Applying theory of one-dimensional diffusion of proteins along DNA, we show that exponential decay of methylation rate correlation with genomic distance is consistent with enzyme processivity. Our results provide quantitative evidence of genome-wide methyltransferase processivity *in vivo*. From the experiment-derived correlations, we estimate that an individual methyl-transferase methylates neighbor CpGs in sequence if they are 36 basepairs apart, on average. But other mechanisms of coupling dominate for inter-CpG distances past ~ 100 basepairs. Our study demonstrates that quantitative insights into enzymatic mechanisms can be obtained from replication-associated, cell-based genome-wide measurements, by combining data-driven statistical analyses with hypothesis-driven mathematical modeling.

## Introduction

DNA methylation is an important epigenetic modification that plays a critical role in development, aging, and cancer, and it is well conserved among most plants, animals and fungi (1, 2). In mammals, DNA methylation occurs predominantly in the cytosine-phosphate-guanine (CpG) dinucleotide context. Across most of the mammalian genome, CpGs occur with low frequency, except for regions called CpG islands (CGIs), which are often associated with promoters (3). Methylated promoters are associated with transcriptional repression, pointing to a role for DNA methylation as a stable and heritable chromatin mark to program alternative gene expression states (4, 5).

The inheritance and maintenance of methylation patterns across cell cycles is important in development and throughout organismal lifespan. Methylation patterns encode information related to gene expression (6, 7), differentiation (8), and genomic imprinting (9). Failure in maintenance and transmission of such patterns can lead to aberrant gene expression, and diseases including cancer (10), developmental abnormalities, and even death (11). The classical model of DNA methylation maintenance introduced the idea that the symmetrical nature of the CpG dinucleotide provides a biomolecular structure whereby DNA methylation could be inherited across a single CpG site by the activity of a, then posited, “maintenance” methyltransferase enzyme (9, 12). The mammalian DNA methyltransferase DNMT1 was subsequently found to serve as the primary maintenance enzyme (13). However, the classical model has been refined in a number of ways based on updated understanding of the biochemical properties and genomic activity of methyltransferases (reviewed in (14)). For instance, a wealth of evidence suggests that the efficiency and specificity of methylation enzymes are not sufficient to support the observed high fidelity of maintenance, within the classical, independent-site model (15–19).

Interdependence, or coupling, of maintenance methylation activities imparted on CpGs located within close proximity can provide some reconciliation between the known biochemistry of methylation reactions and the observed stability of the genomic methylation landscape. Recent findings of preferential recruitment of DNMT1 (20), and faster maintenance methylation rates (21), at sites with more neighboring hemimethylated CpGs are clearly at odds with an independent-CpG-site model. CpG interdependence has been suggested to occur via various molecular mechanisms, including DNMT1 processivity (22–24) (in which an enzyme can methylate multiple neighboring CpGs on nascent DNA sequentially) and cooperative interactions, e.g., with UHRF1, which localizes, and in turn helps recruit DNMT1 molecules, to hemi-methylated CpGs (25, 26). Mathematical modeling has also suggested the importance of CpG interdependence, also called collaboration, both in maintenance and *de novo* methylation, for long-term collective stability of methylated and unmethylated genomic regions (19, 27–31).

A quantitative and mechanistic understanding of CpG interdependence during maintenance methylation *in vivo* is lacking. The genomic lengthscales over which CpG coupling occurs are not well understood. It is not yet known to what extent processivity, versus other mechanisms of CpG interdependence, influences dynamics of maintenance methylation. Nor is it yet well-understood how local genomic context influences these mechanisms *in vivo*. In this study, we address these questions by elucidating CpG-coupled-dynamics in maintenance methylation by use of statistical inference, bioinformatics, and stochastic modeling. We leverage experiments that measured methylation status of nascent-strand CpGs across post-replication timescales, genome-wide (32). From these data, we infer how the rates with which individual CpGs acquire methylation, post-replication, are correlated on nearby sites in different regional contexts. Using stochastic models and theory of proteins diffusing along DNA, we demonstrate that the rate correlation as a function of genomic distance provides a mechanistic fingerprint for post-replication enzymatic processes. Our novel method provides the first direct evidence for genome-wide methyltransferase processivity in cells. Comparing simulations to data allows us to extract quantitative insights from data, including the relative strengths of processive versus non-processive coupling mechanisms in different genomic regions, and the length of processive steps.

## RESULTS

### DNA methylation *rate* and *state* on neighboring CpGs are correlated to different extents

An analysis of cytosine methylation within newly replicated DNA over time (via Repli-BS) revealed that some genomic loci exhibit a pronounced lag in methylation maintenance (32). In our previous work, we developed a statistical inference procedure to obtain single-CpG post-replication maintenance methylation rates (here denoted “remethylation rates”) from Repli-BS data (33), which further supported the variability of remethylation kinetics across the genome. Other recent studies found variable kinetics of maintenance methylation in different genomic contexts (21) and of combined methylating/demethylating reactions (34). However, these studies did not investigate whether or how methylation rates vary locally within regions or among neighboring clusters of CpGs.

In this paper, we focus on local correlation of methylation kinetics, obtained from Repli-BS data, and analyze the data-derived correlation using biophysical models of enzymatic methylation reactions. The data-inferred kinetic parameters quantify the rate of accumulation of methylation at each CpG site across hESCs in the measurement set over the experimental timecourse of 0-16 hours post-replication. We correlate single-CpG remethylation rate constants (denoted *k_i_*, for the rate at the *i*th CpG, obtained by Maximum Likelihood Estimation, see Methods) on pairs of CpGs, as a function of the genomic distance between them. As such, this postreplication methylation *rate correlation* quantifies the extent to which CpG pairs experience similar (fast or slow) kinetics. DNA methylation levels (also hereon denoted methylation “states”, or the fraction of cells exhibiting methylation at an individual CpG site) on neighboring CpGs are also correlated (30, 35). This methylation *state correlation*, as a function of genomic distance in basepairs, reflects information such as the size of persistently methylated (or unmethylated) domains. As such, it reflects the generally static methylation landscape in a given cell type.

We reasoned that correlation of post-replication methylation rates on individual CpGs could reveal details of enzyme kinetics in cells, and thereby yield additional information beyond that contained in methylation state correlation. To this end, we first computed correlation from the two different data modalities ((i) remethylation rate derived from Repli-BS and (ii) methylation state derived from WGBS) to relatively short (< 1 kilobasepair) distances. We compared them across different genomic contexts, by filtering CpG pairs by genomic context in addition to their inter-CpG distance (Figure 1a). Genomic regional contexts were based on genomic annotations acquired from the UCSC genome table browser and included features such as 3’UTRs, enhancers, etc. (see Methods).

**Fig. 1.**
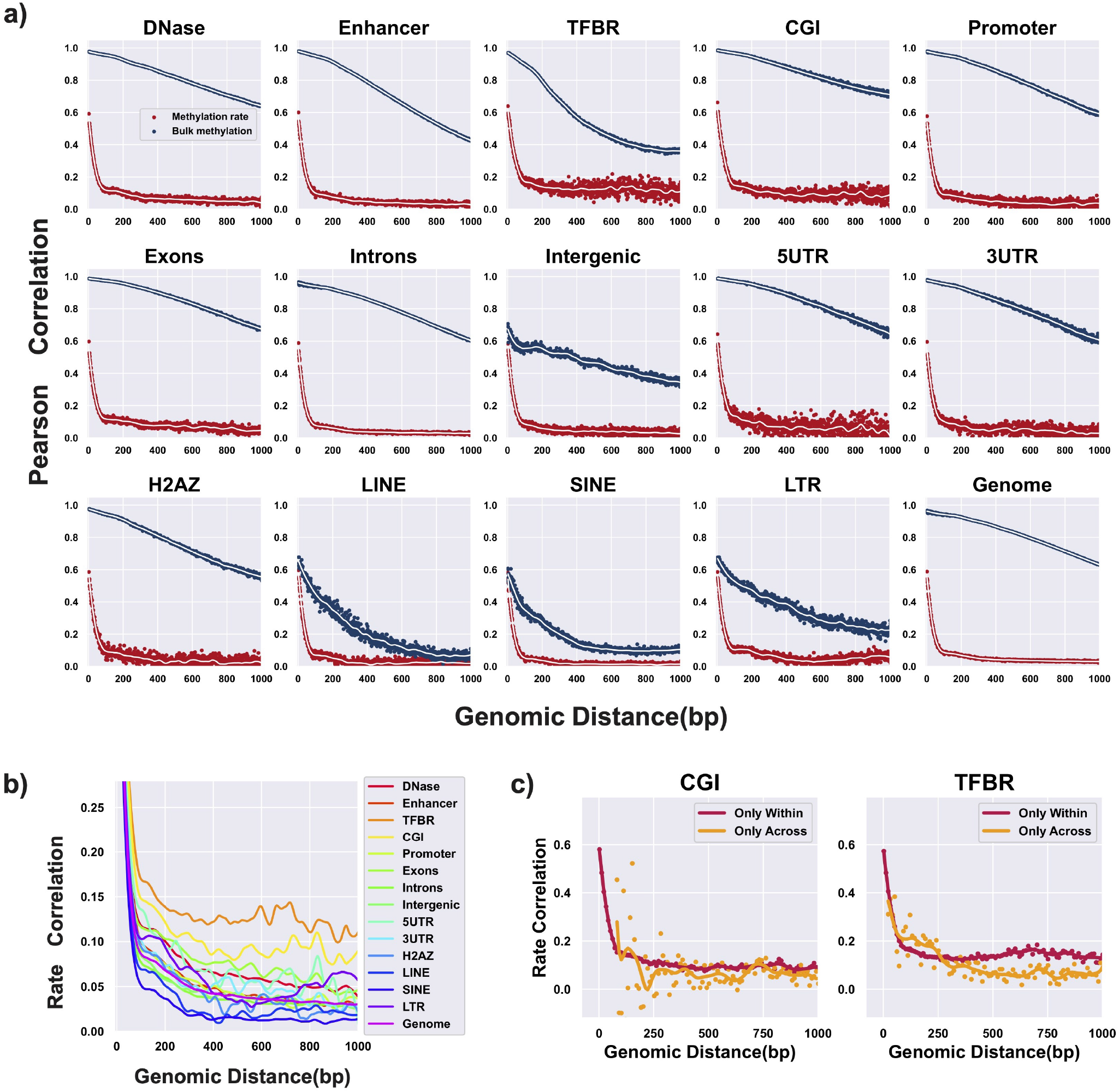
Methylation states and post-replication remethylation rates are correlated on neighboring CpGs, and correlation varies with genomic regional context. **(a)** Correlation of remethylation rates (red, y-axis) and correlation of percent methylation (i.e., bulk methylation state, blue, y-axis) of nearby CpGs at given genomic distances (x-axis), separated in panels by genomic regional context. The remethylation rates for individual CpG sites were inferred from experimental Repli-BS-seq data, and the bulk methylation was obtained from separate WGBS data from the same cell line (HUES64). Dots: raw rate correlation in base pair resolution, lines: smoothed correlation curve by LOESS(LOcally Estimated Scatterplot Smoothing,(36)) with 100 bp span. **(b)** Correlation of remethylation rates with curves from different genomic regional contexts overlapped and y-axis zoomed. Only the LOESS smoothed curves are shown for clarity. **(c)** Rate correlation functions computed retaining only pairs of sites identified either as within the same localized region (red) or not within the same localized region (orange).

We observe both common and distinct features in the correlation functions, when comparing different genomic regional contexts for both data modalities (i.e., rate-correlations and state-correlations). Common to all regions, and to the genomic average with no region-filtering, remethylation rate-correlation decreases rapidly with distance until approximately 100 bp (red curves in Figure 1a). After 100 bp, a slower decay is evident in all regions. However, this steep decrease in correlation and rapid change in decay is not observed in correlation of methylation state (navy curves), which generally shows a slower (persisting to *>*1 kb), smooth decay, albeit with wide variation in decay-lengths between regions. The existence of such a pronounced difference between methylation state and rate correlation, persistent in all genomic regions, suggests that the methylation rate correlations are not determined by the methylation landscape itself, and raises the question of what dynamic processes determine rate correlations.

### Post-replication remethylation rates are correlated on neighboring CpGs, and correlation varies with genomic regional context

Some features of the correlation functions vary between the genomic regions studied. We quantify the average magnitude of short-range correlation by the mean correlation to 1 kbp. These magnitudes are highly variable across genomic contexts and across modalities. Genome-wide, the magnitude of rate-correlation is low (0.06) compared to that of state-correlation (0.82). Across regions, CGIs (CpG Islands) show the highest magnitude of state-correlation (0.85) while transcription factor binding regions (TFBR) shows the highest magnitude of rate-correlation (0.15). By contrast, the lowest average magnitude of state-correlation and of rate-correlation are both found within SINEs (Short Interspersed Nuclear Elements) regions with 0.18 and 0.035 correlation, respectively. We hereafter refer to these differences in remethylation rate- and state-correlations according to genomic context as “genomic region specificity”.

In general, the results for methylation state indicates that methylation on CpGs within 1kb are highly correlated genome-wide. Exceptions are within SINEs and LINEs (Long Interspersed Nuclear Elements), where state-correlation has significantly decayed by 1 kbp.

Although the methylation rate correlations show an apparently uniform sub-100-bp decay across regions, differences are visible in their longer-lengthscale (>100 bp) decay profiles (Figure 1b). In particular, CGIs and TFBRs (Transcription Factor Binding Regions) show the strongest longdistance correlation, persisting near or above 0.1 past 1 kbp. We further separated the analyzed CpGs from CGIs and TF-BRs into “within” and “across” region pairs (1c). “Within” correlation is computed for CpGs that are within one contiguous region (as defined by the filtering protocol), whereas “across” correlation only retains pairs of sites that are assigned to non-contiguous regions. The resultant rate correlation functions indicate generally stronger contribution of within-versus across-region correlation (note that the number of datapoints for “across” correlations is lower for short distances, since nearby CpGs are more likely to be in contiguous regions). In all, these results indicate different magnitudes and decay-lengths of inter-CpG coupling in maintenance methylation. In particular, they point to more pronounced CpG collectivity of maintenance kinetics in contiguous CGIs and TFBRs. That is, neighboring CpGs in CGIs and TFBRs tend to have more correlated remethylation rates and thus more similar methylation across post-replication time, as compared to other regions. However, all regions showed correlated rates to some degree.

We note several additional features in the correlation functions, including apparent periodicity (e.g., in state-correlation, Intergenic region, Figure 1a) and shoulders or local peaks (e.g., in several of the rate-correlation curves in Figure 1b). Some of these features appear consistent with nucleosome positioning, which could indicate coupling between maintenance methylation processes and/or the methylation landscape with nucleosomes. A detailed investigation of all of these features is outside the scope of this work; we focus instead on the broad features of the methylation rate correlation, namely the sub-100 bp rapid decay and region-specific, > 100bp slower decay.

### Regional correlation of DNA methylation maintenance kinetics is increased with CpG density and chromatin accessibility, and decreased with higher bulk methylation levels

To investigate the factors associated with the observed region specificity, the association between remethylation rate correlation and other local genomic characteristics were examined. For each individual CpG in the dataset, a measure of the local chromatin accessibility is collected from DNaseI hypersensitivity data in ENCODE/OpenChrom (see Materials and Methods). The local CpG density surrounding a given site is calculated by the number of neighboring CpGs within a 500bp window. “Bulk” methylation refers to measurement of DNA methylation in human embryonic stem cells(HUES64) measured by Whole Genome Bisulfite Sequencing(WGBS). We plotted the magnitude of the remethylation rate correlation (0-1000 bp) for each region versus average chromatin accessibility (e.g., DNAse hypersensitivity enrichment) (Figure 2a), local CpG density (Figure 2b), and WGBS (“bulk”) methylation percentage (Figure 2c). In all cases, a linear correlation was found between the magnitude of remethylation rate correlation and these three factors; rate correlation is positively correlated with DNase level and CpG density (albeit weakly), while it is inversely correlated with bulk methylation level. These results suggest that the genomic regional differences observed in rate correlation in Figure 1 may be driven globally by variation in chromatin accessibility, CpG density, and background methylation landspace.

**Fig. 2.**
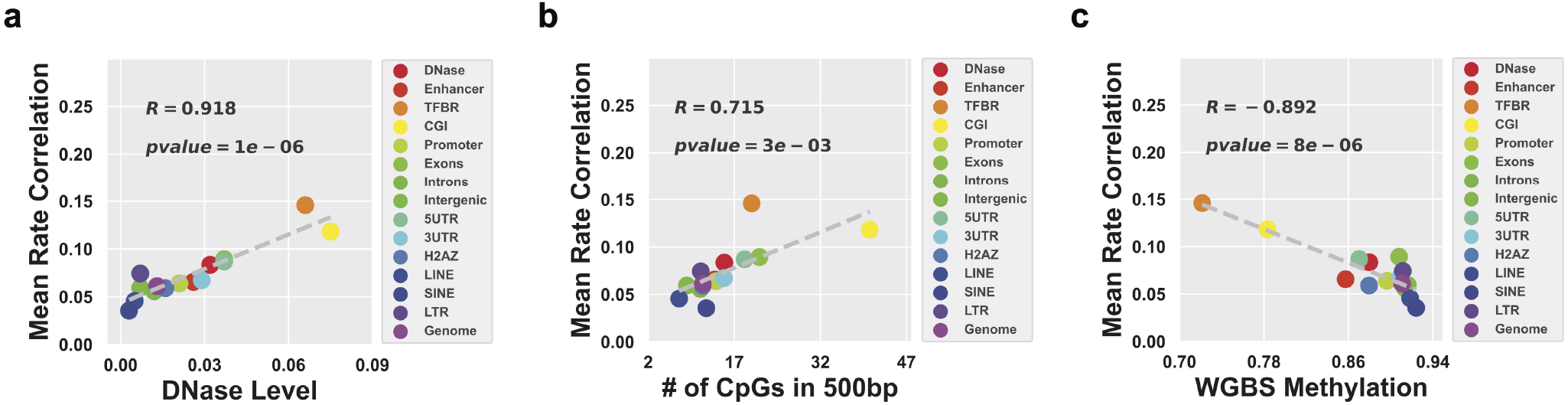
DNA methylation maintenance rates show higher local correlation in genomic regions with higher chromatin accessibility, CpG density and lower bulk methylation levels. Mean remethylation rate correlation by region, plotted versus other quantified, localized genomic measurements from independent measurements: **(a)** Magnitude of mean remethylation rate correlation (equal to the average over rate correlation for all inter-CpG distance < 1 kbp (i.e., integers from 2-1000) in the given region) versus mean regional chromatin accessibility, as quantified by DNase level; **(b)** Versus mean local CpG density (mean number of neighboring CpGs within a 500bp window); **(c)** Versus mean WGBS (“bulk”) methylation level. Note that the datasets are site-matched, so the analysis is restricted to sites that tend to have intermediate to high methylation, since these are the sites for which remethylation rates are available.

### Local methylation correlation varies across post-replication time and shows persistent region specificity

In order to investigate the origins and regional differences of methylation correlation, we focus on three representative genomic regions characterized by high (CGI), medium (Enhancer) and low (SINE) methylation state correlation. In figure 3, we plot three types of correlation functions for these three regions: in addition to bulk WGBS correlation and rate correlation, we also plot correlation of methylation state in “nascent” DNA, which contains a subset of measurements from the full temporal Repli-BS dataset, corresponding to the 0-hour timepoint of the original pulse-chase experiment(32). Thus, “nascent” here refers to methylation readout ≤ 1 hour post-replication.

**Fig. 3.**
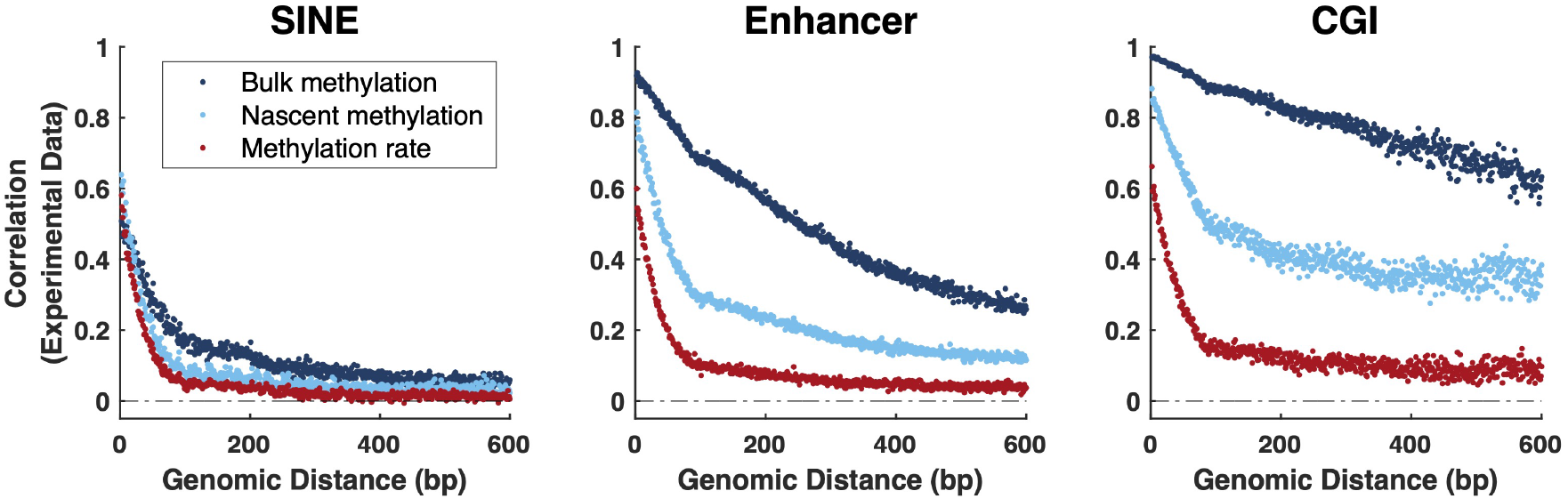
DNA methylation correlation increases over post-replication time and shows persistent region specificity across different data modalities. Data from three representative regions and three methods of extracting methylation correlation; all three curves in each panel are derived from the same set of CpGs. Dark blue: correlation of methylation state of neighboring CpGs from Whole Genome Bisulfite Sequencing Experiments (WGBS, “Bulk”). Light blue: Correlation of methylation state of neighboring CpGs from WGBS on nascent DNA from Repli-BS-seq experiments (i.e., < 1 hour post-replication) (“Nascent”). Red: Correlation of remethylation rates inferred from Repli-BS-seq experiments. Data for rate-correlation (red) is identical to that of Figure 1a. Data for bulk methylation (WGBS) is also derived from the same dataset as Figure 1a, but filtered to retain only those CpG sites for which a rate was available (determined by per-site read-depths, see Methods). Note that the nascent (light blue) and rate (red) curves are not from independent datasets, since the nascent data is a subset of the total Repli-BS-seq dataset from which the rates were inferred. Blue Pearson correlation functions are computed from raw data, whereas the red correlation function is computed on inferred rate parameters from the MLE pipeline.

Note that the red rate and dark blue state correlations are derived from the same data as in Figure 1, but in contrast to Figure 1, the curves in each panel of Figure 3 are site matched. In practice, remethylation rates are available for only a subset of CpGs, as compared to WGBS measurements. This is because the rate constant is undefined where no methylation is measured, and is often unidentifiable when methylation is very low. Thus, there is significant overlap between the sites for which rates are not available and sites for which WGBS percentages are 0 or near 0. We reasoned that some of the methylation state correlation in Figure 1 could arise from the bimodal nature of methylated and unmethylated regions. For a more direct comparison to rate correlation on a site-specific basis, we thus filtered to a common set of sites, which in practice retains mostly sites with intermediate to high methylation in WGBS. After filtering to these common sites, we indeed observe some decrease in state correlation (dark blue curves in Figure 1a versus 3), but rate correlation remained persistently lower than state correlation in all regions.

We observe that methylation state correlation in nascent DNA is generally lower than that in bulk DNA, suggesting that state correlation increases over post-replication time. (Bulk WGBS measurements reflect temporal variability from cells in various stages of the cell cycle and from differences in replication-timing across the genome (meaning it largely contains matured DNA strands), whereas the nascent data in principle captures reads within one hour post-replication (32)). In all studied regions, the nascent methylation correlation is intermediate between rate and bulk state correlation (with the exception of very short distances in SINEs).

The trend in genomic region specificity of methylation correlation is persistent across the three types of correlation functions. This in turn suggests that the region specificity is persistent across post-replication time. For example, CGIs consistently show the highest correlation compared to the other regions in rate, bulk methylation, and nascent methylation. Conversely, SINE consistently shows the lowest correlation. These results suggest that the processes that govern coupling (or interdependence) of methylation among neighboring CpGs differ depending on the genomic regional context, and that these processes are region-specific already at early post-replication timepoints. Additionally, the distinct shapes and magnitudes of the CpG-neighbor correlations across time suggest that different processes control CpG-neighbor-interactions at early versus late post-replication times.

We hypothesize that the three correlation modalities can be interpreted as follows: rate correlation reflects the dynamic mechanisms of maintenance methylation, thus shedding light on early post-replication-time processes. In contrast, bulk methylation state correlation largely reflects the steady-state methylation landscape, i.e. reflecting the balance among methyl-reading/writing/erasing processes operating across post-replication time to regulate the methylation landscape, but largely reflecting the stable methylation landscape of a given cell type. Nascent methylation state reflects a mixture of the two, as the experiment “captures” CpGs in transit between their state immediately (up to one hour) postreplication and steady state. In the following sections, we test this hypothesis by use of computer simulations and model-guided data analysis.

### Region-specific stochastic simulations of post-replication maintenance methylation

In order to gain further biological insight from the experiment-derived methylation correlation functions, we perform region-specific stochastic simulations of maintenance methylation (Figure 4), and use these simulations to generate synthetic data analogous to the various experimental bisulfite sequencing data modalities. From these synthetic data, we compute regional correlation functions and compare to those derived from experiments. Briefly (see Methods), the simulations track nascent-strand methylation status of stretches of sequentially positioned CpGs, numbering on the order of tens of thousands. In contrast to mathematical models that treat the interplay of *de novo*, maintenance, and demethylating reactions, e.g. (19), we apply minimal models of single-site-resolution DNMT1-mediated methylation on post-replication timescales. Each stochastic simulation tracks the binary (methylated or not methylated) status of the “nascent-strand” CpGs. At the start of the simulation (representing exactly time 0 with respect to DNA replication at that site-i.e., the time of nucleotide addition), all nascent CpGs are assumed to be unmethylated. The presence or absence of methylation on cytosine bases on the opposing parental-strand at time 0 is determined probabilistically from a data-derived regional methylation landscape that acts as the simulation input. If the parental cytosine is methylated at time 0, then the CpG is considered hemimethylated and the nascent cytosine is assumed to be a target for DNMT1-catalyzed methylation, and it will acquire methylation stochastically at some post-replication timepoint, according to the chemical reaction kinetics encoded in the model. If the parental cytosine is unmethylated at time 0, then DNMT1 does not target the nascent cytosine for methylation, and the site will remain fully unmethylated. In this way, the model tracks only unidirectional maintenance methylation, and does not include active demethylation reactions. It also does not account for any de novo methylation activity. The simulation tracks post-replication timescales (following experiments, to approximately 16 hours), up to but not including subsequent replication events.

**Fig. 4.**
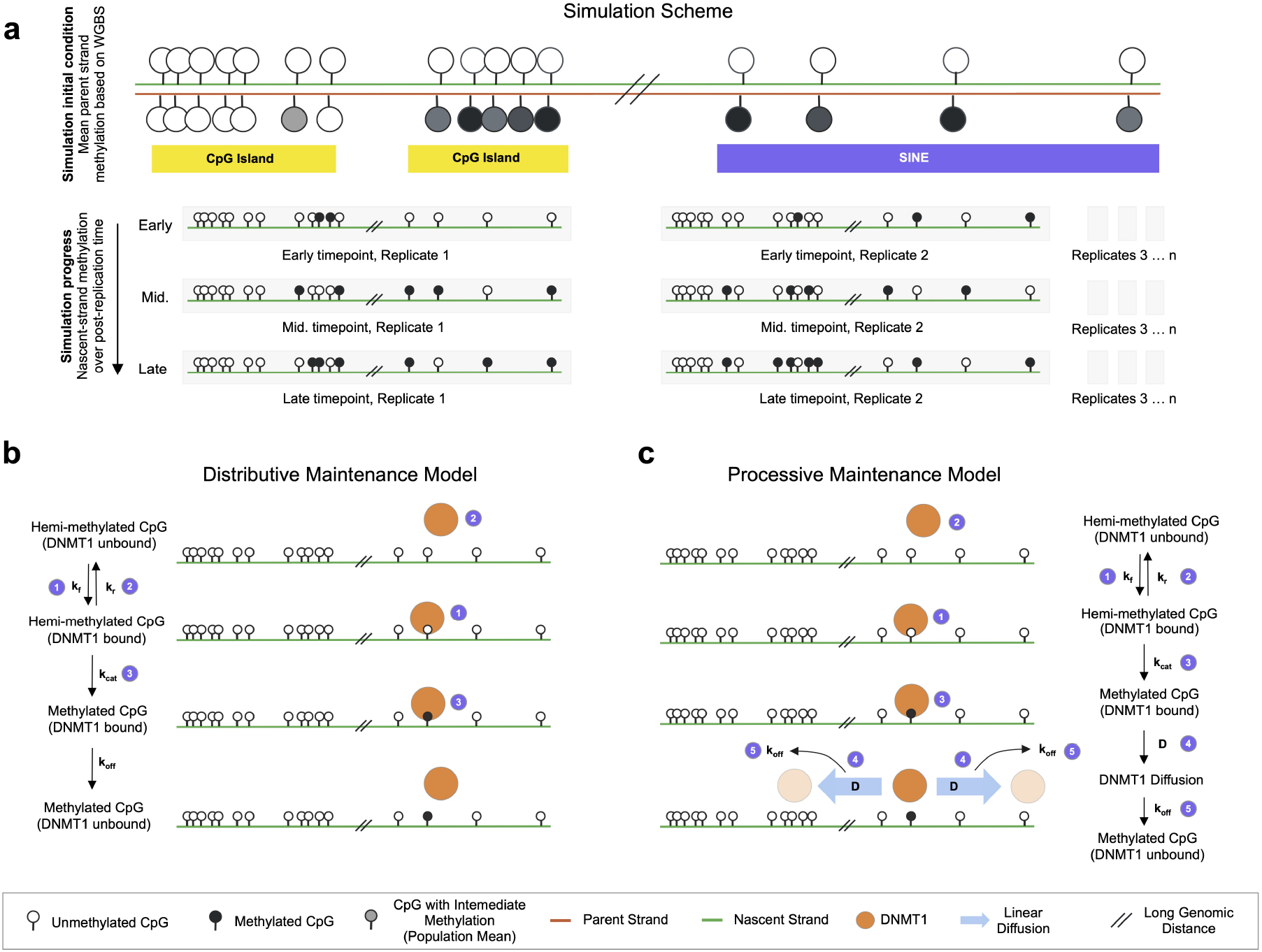
Schematic of stochastic simulation of DNA methylation maintenance in different genomic regions, according to either Distributive or Processive mechanisms. **(a)** Each simulation models a strand of *N* CpGs (*N* = 25000 to 75000) where the CpG positions and the parent-strand methylation at the initial condition are taken from WGBS measurements in a given genomic region (e.g., CGI or SINE). Multiple strand replicates are simulated over time. Immediate-post-replication DNA is assumed to be unmethylated on the nascent strand at all sites, and the binary methylation status of the parent strand sites are sampled probabilistically from the input methylation landscape (mean methylation level in WGBS). During the simulations, DNMT1 targets hemimethylated CpGs for methylation, according to either a Distributive **(b)** or Processive **(c)** mechanism. The Distributive mechanism assumes that the enzyme binds to each hemi-methylated CpG independently. After catalyzing methylation, the enzyme immediately unbinds from DNA (we assume *k*_off_ ≫ *k*_cat_, such that methylation and unbinding are treated as a single reaction). To reach a subsequent CpG, the enzyme must independently rebind with rate *k_f_*. The Processive mechanism assumes that, after catalyzing methylation at a CpG, DNMT1 can remain bound to DNA and reach nearby hemimethylated CpGs by linear diffusion along DNA.

Region-specificity is encoded at the start of the simulation in two ways: (1) the CpG positions and (2) the local methylation landscape, meaning: the probability of the nascent CpG to be a target for DNTM1-catalyzed methyl-addition due to the presence of methylation on the parental strand CpG dinucleotide. Both of these quantities are derived from experimental WGBS data with a regional filter to retain only CpGs in the desired region. Thus, in simulating a CGI region, the *i*th simulated CpG (*i* ∈ [1…*N*]) has a genomic position *x_i_* and a probability *f_i_* to be targeted for methylation. We obtain both *x_i_* and *f_i_* from WGBS data from hESCs, where *x_i_* is the integer site-ID for the cytosine (which is identified as being located within a CGI) and *f_i_* ∈ [0, 1] is taken to be equal to the measured methylation fraction at that site. For example, if a given CpG in the dataset has a WGBS-measured methylation fraction of 0.8, then the model assigns the parental cytosine to be methylated at the start of the simulation with a probability equal to 0.8. Strands are simulated in replicate. With sufficient replicates, the simulation eventually recapitulates the experiment-derived input methylation landscape, if it is run for a period of time that is sufficiently long. Note that the WGBS-data-derived landscape likely reflects some degree of replication-associated temporal variability (32), rather than a true steady-state. Nevertheless, use of the WGBS-background-methylation landscape as the simulation input allows us to encode realistic region-specific differences in CpG densities and qualitative differences in bulk methylation levels.

### Rate correlation provides a mechanistic fingerprint for enzyme kinetics

We use simulations to generate synthetic data mimicking the various bisulfite sequencing datatypes (rates from Repli-BS, nascent methylation from Repli-BS, and bulk methylation state from WGBS). We then compute the correlation functions for the synthetic data. Figure 5 shows simulation-derived correlation functions for three representative genomic regions from Chromosome 1 for two mechanistic models termed Distributive and Processive. Briefly, DNMT1 binds to nascent CpGs and catalyzes the addition of a methyl group. In the Distributive model (Figure 4b), the enzyme unbinds after the catalytic step, and must independently rebind to an available hemi-methylated CpG in order to catalyze a subsequent methyl addition. In the Processive model (Figure 4c), the enzyme can remain bound to DNA after methylating a CpG, and can travel along DNA (via a one dimensional diffusive random walk) to reach neighboring hemi-methylated CpGs and again catalyze methylation. The random walk occurs with with 1D diffusion coefficient, *D*, and the enzyme potentially unbinds before reaching its target with rate *k_off_*. The kinetic parameters for both models are given in the Supplement.

**Fig. 5.**
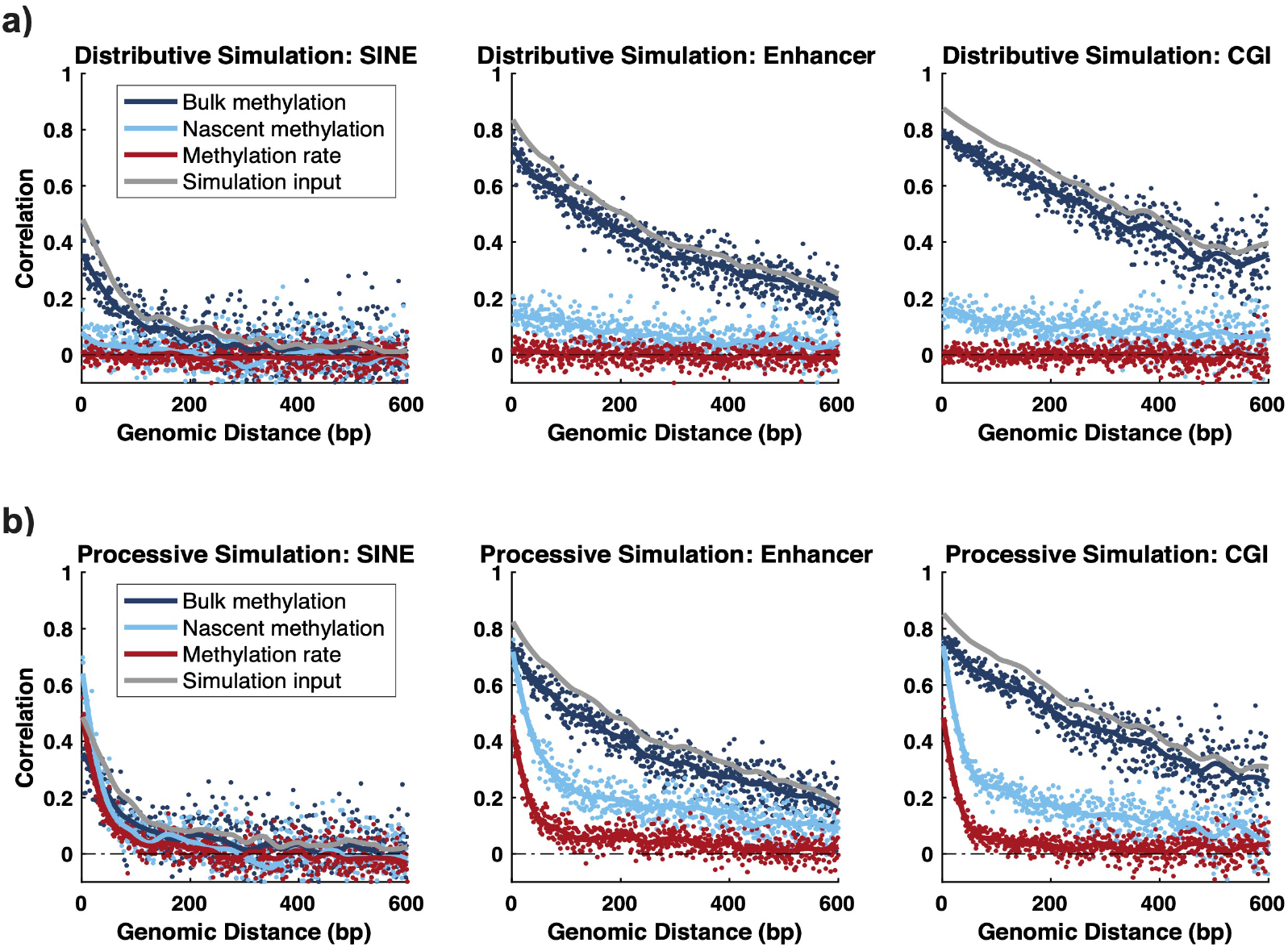
In simulations, rate correlation but not state correlation depends on enzymatic mechanism; only Processive model displays nonzero rate correlation. Simulated correlation functions for three representative regions (SINE, Ehnhancer, CGI) and using the Distributive (top) or Processive (bottom) mechanism. Simulations are performed using experimental regional methylation landscapes for SINE, Enhancer, and CGI as initial condition inputs (grey curves). The simulations provide synthetic data corresponding to each of the three experimental data modalities, as in Figure 3. Synthetic data are processed in the same way as the experimental data to compute the inferred remethylation rates and Pearson correlation functions. Both models recapitulate temporal trends in rate correlation seen in experiments, Figure 3 (correlation in rate < nascent state < bulk state). Only the Processive model captures nonzero rate correlation sub-100-bp; neither model captures low, persistent correlation > 100 bp observed in Figure 3.

In both Distributive and Processive mechanisms, the simulated bulk methylation correlation approaches the input methylation landscape in the three regions. This is expected, because the input landscape dictates parental strand methylation in the model, and the model assumes that as time progresses, the nascent strand methylation will ultimately match the parental strand input (i.e., it assumes perfect maintenance in the long-time limit). In contrast, the shapes of remethylation rate correlation are distinct from those of the input methylation landscape, and are dependent on the model mechanism. In all simulated regions, nascent correlation is intermediate between bulk state and rate, in agreement with experiments. The Distributive model produces no correlation in remethylation rate, in agreement with our previous results (33). In contrast, the Processive model produces a nonzero rate correlation that appears to qualitatively reproduce the experimentally observed rapid decay of correlation with genomic distance up to approximately 100 bp. Neither model reproduces the low but persistent correlation visible at distances greater than 100 bp in CGI and Enhancer in Figure 3.

The simulated correlation functions provide a possible explanation for the discrepancy between methylation state and rate correlations observed from experiments (Figures 1 and 3). Namely, they suggest that the methylation rate correlation shape is dictated by enzymatic maintenance mechanisms, independent of the background methylation landscape. That is, simulations support that mechanistic insights on maintenance methylation can be derived using the rate correlation, since it effectively separates correlations introduced by early post-replication-time processes from those operating at longer timescales (and which dominate the bulk WGBS correlation). Thus, the rate correlation is a useful new quantity, which can be used to distinguish between hypothesized mechanisms. The simulations also demonstrate that, while the Processive mechanism partially recapitulates experiment-derived rate correlations, it cannot explain the longer-distance, slow rate correlation decay in CGI.

### Rate correlation in Processive model decays exponentially, dependent on 1D diffusion and unbinding rates

We used simulations and theoretical modeling to determine whether quantitative insights could be derived from the experimental rate correlations. We first investigated how rate correlations arising from the Processive mechanism depend on model parameters. In an idealized theoretical model, we find that the rate correlations are related to the diffusion constant *D* and the unbinding rate, *k_off_*, as:

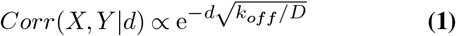

where *X*, *Y* are pairs of remethylation rates on sites that are distance *d* apart (along the DNA strand). Equation 1 can be understood as follows:

Consider two target sites labeled *i* and *j*, located on a 1D strand of DNA, that are located some distance *d* away. Let *τ_i_* be the time (post-replication) at which site *i* acquires methylation (similarly, *τ_j_*). Assume that in a particular realization of the stochastic process, two possible events can occur:

1. Each of the sites is independently methylated, in which case *τ_i_* and *τ_j_* are independent samples from some common distribution, denoted 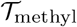. 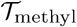 reflects the distribution of times required for all biochemical steps in the site acquiring methylation, e.g., for DNMT1 to bind DNA from the nucleosol, form a complex with the CpG site, and successfully catalyze methylation.
2. Alternatively, one of the sites could be methylated by an enzyme that has reached it by diffusion after first acting on the other site. In this case, assuming (without loss of generality) that site *i* is the first, then *τ_i_* is sampled from 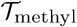 and *τ_j_* = *τ_i_* + *τ_D_*, where *τ_D_* is sampled from random variable 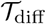. 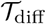 reflects the distribution of times for DNMT1 to diffuse the distance *d* to reach the neighbor site, form a complex, and successfully catalyze methylation.

The likelihood of event 1 or 2 above taking place depends in part on the probability that the enzyme can remain bound to DNA and diffuse a distance *d* before unbinding. In the limit of 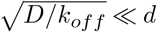, then event 1 always occurs, and

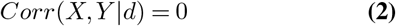

where *X*, *Y* now denotes pairs of methylation times at sites with intervening distance *d*. The correlation is zero because *τ_i_* and *τ_j_* are assumed to be i.i.d. random variables. Conversely, in the limit of 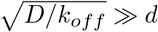, then event 2 always occurs, and

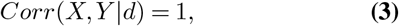

(in the limit of *τ_D_* ≪ *τ_i_*). Assuming that the enzyme need only *reach* the next target site by diffusion, in which case the catalytic step occurs subsequently with probability 1, then the correlation is equal to the probability of event 2 occuring, i.e.

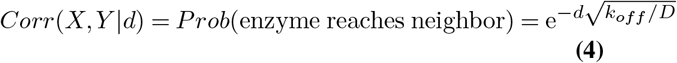

where the exponential dependence on linear distance, with decay rate 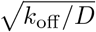, is obtained from a 1D analytical diffusion model with parameters *k*_off_, *D*, on a semi-infinite domain with an absorbing state at the target site at distance *d* from the starting point. Such models have been used previously for theory of transcription factor searching on DNA (37). In the full Processive model, the enzyme does not always successfully methylate the target upon reaching it, leading to the proportional relationship of Equation 1. The complete analytical results are presented in the Supplement.

We find a good match between the above analytical theory and the stochastic simulations (Figure S1). We performed regional simulations for varying values of *D/k_off_*, and then processed the simulated data through the MLE inference pipeline and computed the correlation function. We then fit the correlation functions to a single exponential decay, and observe a good match between the fitted decay constant and the theoretically predicted value of 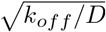. These results demonstrate that the predicted exponential correlation holds, despite potentially complicating factors in the stochastic model (e.g., the finite time required for diffusion, many CpG sites and multiple enzymes acting simultaneously on one strand, etc.), and uncertainty introduced by the MLE-fiting pipeline. All in all, these results support the finding that processivity, in a linear diffusion model, is consistent with exponential decay of rate correlations obtained from Repli-BS, and that the decay constant can be interpreted as 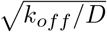.

### Mathematical model for diffusion-based and region-based components of methylation rate correlation

The Processive model can explain exponential decay of rate correlation, but cannot by itself explain the significant correlation observed past 100 bp in strongly correlated regions such as CGI (Figure 3). Nor can it explain the association of this correlation to other regional genomic characteristics (bulk methylation levels, CpG density, chromatin accessibility). Based on the observations in Fig 2, we reasoned that this additional rate correlation can be understood to result from correlated methylation times on neighboring sites due to a variety of regional features, that we describe as

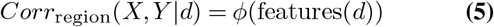

and refer to as “region-based correlation”. That is, this component of the correlation function can be expressed as a function *ϕ*, which depends somehow on local features, e.g., related to the chromatin landscape. These features themselves held some distance-dependent correlation. Consideration of additional correlating factors (beyond diffusion) equates to a revision of the idealized theoretical model above. Now, in event 1, if the neighboring site is not reached by diffusion, but rather the two sites are methylated in separate events, their times *τ_i_* and *τ_j_* are nevertheless correlated in a way that depends not only on their distance apart, but also on various other features of their location within the genome. We do not propose a mechanistic model for this additional correlation, but label it as *ϕ*, and estimate it for each region, based on the data. Given these two contributions, which we term diffusion-dependent (denoted *θ*) and region-dependent (*ϕ*), the total correlation from both contributions is given by:

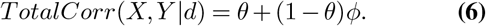

where the above equation can be understood by probabilistic arguments, i.e., the probability that a neighbor is reached by diffusion is *θ* (given by Eq.1); additional correlation *ϕ* is only present when the neighbor is *not* reached by diffusion, with probability 1 – *θ*. This mathematical decomposition of the rate correlation is shown schematically in Figure 6a. The model of equations 1, 5, and 6 makes a prediction: that if *ϕ* (the component of correlation due to genomic regional features) can be estimated from the data, then the remaining correlation (*θ*) should decay exponentially with distance, and should not depend on local genomic context.

**Fig. 6.**
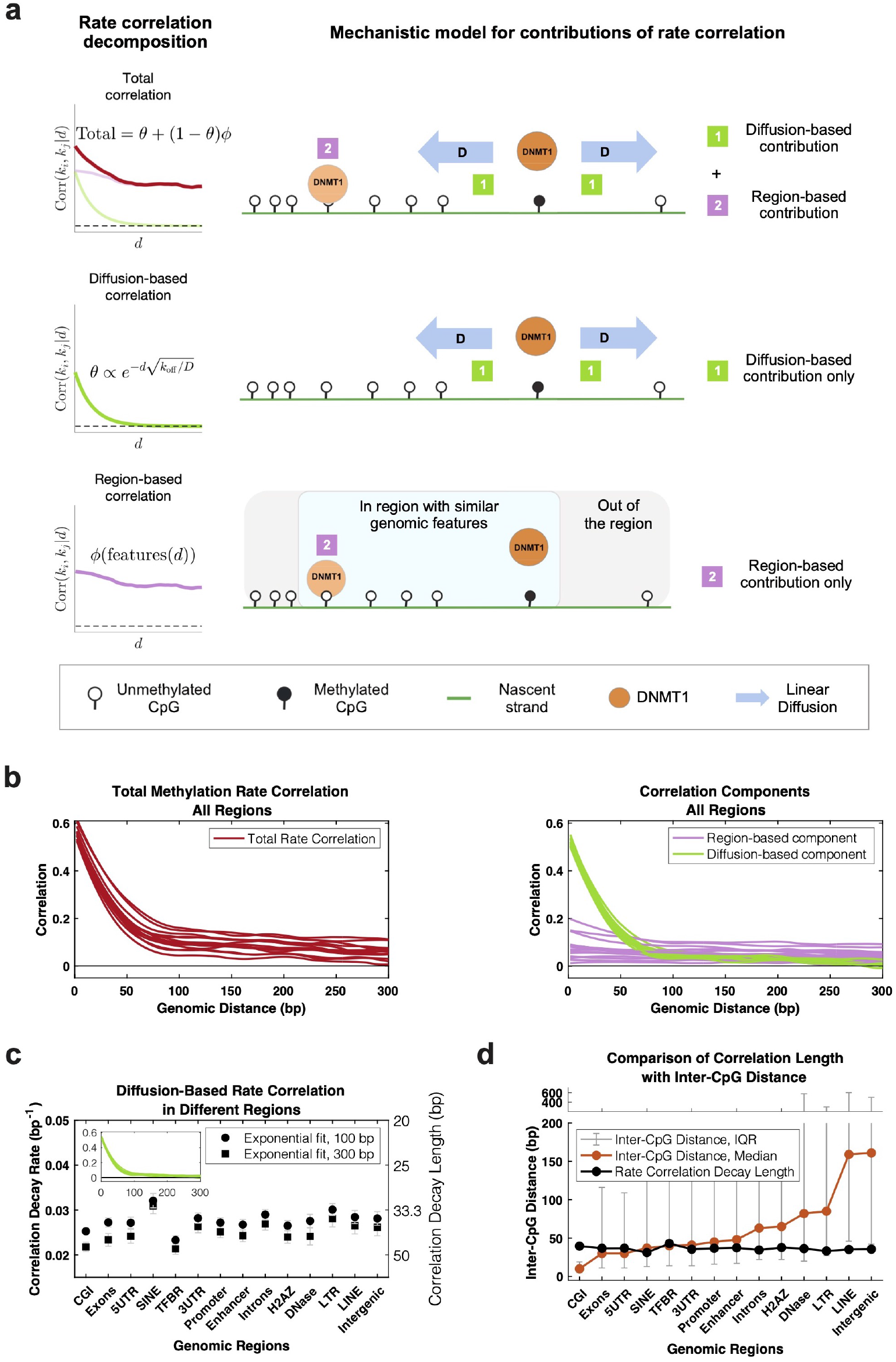
Decomposition of experimental rate correlation into processive and nonprocessive components allows estimation of DNMT1 diffusion parameters from data. **(a)** Mathematical model of mechanistic contributions to rate correlation predicts that the total rate correlation (red curve) arises from two types of processes, termed diffusion-based (i.e., processive) and region-based (i.e., nonprocessive). The diffusion-based component (green curve) is due to an individual enzyme at CpG_*i*_ reaching nearby target CpG_*j*_ by linear diffusion, and shows exponential decay with genomic distance *d*. The region-based component (pink curve) is attributed to any other sources of correlation in remethylation rates, e.g., when two different enzyme molecules reach their targets (CpG_*i*_ and CpG_*j*_) with correlated arrival times, and does not have an analytic expression. **(b)** Rate correlation of all regions from experimental data are plotted in red in left panel (same red curves as Figures 1a), and the corresponding decomposed components (see Methods) are plotted in right panel, with diffusion-based components in green and region-based components in pink. **(c)** Single-exponential fit decay constants for the green curves (*θ*), using either a 100 bp or 300 bp fitting window. The curves are well-fit in each region by a single exponential decay function, supporting attribution of this correlation component to the processive mechanism (i.e., to diffusion). The decay constants are relatively uniform across regions, supporting common diffusive dynamics across the genome. The average decay constant, 0.028bp^-1^ provides an estimate of 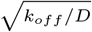, and corresponds to an average distance traveled by DNMT1 to a nearby target CpG of 36 basepairs (values from 100-bp-window). **(d)** Comparison of the inter-CpG distances and extracted rate correlation decay length (from *θ*) of each genomic region, showing that the exponential correlation decay length is insensitive to inter-CpG distance.

### Separation of experimental remethylation rate correlation functions into diffusion-dependent and region-dependent components quantifies genome-wide processivity of DNMT1 *in vivo*

We developed an approach to estimate *ϕ* as follows. The total experiment-derived correlation functions are computed from a list of cytosine positions and their inferred remethylation rates. Denote 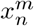 as the position of site *n*, identified as belonging to region *m* (where *m* is one of the fourteen regions of Figure 1, and *n* ∈ [1, *N*] given that data is available for N sites in a region of interest). Denote the remethylation rate as 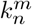, giving a list of pairs {*x_n_*, *k_n_*}_*m*_ for each region, from which the correlation is computed. We also have additional genomic feature data: let 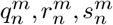 denote the measured bulk methylation level, the local CpG density, and the chromatin accessibility acquired from independent datasets (see Methods). We then perform unsupervised k-means clustering on the features {*q*, *r*, *s*}_*m*_ to obtain clusters of sites that are (i) previously assigned as belonging to the same type of genomic region *m*, and (ii) share more fine-grained similarity in terms of their bulk methylation level, CpG density, and chromatin accessibility. We then randomly shuffle the nucleotide positions within each subcluster, and denote these shuffled positions as 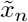. We now have a new list, 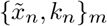, where the true nucleotide positions have been randomized, but their reassigned position is still similar to the true position in terms of the features {*q*, *r*, *s*}_*m*_.

We recalculate the correlation functions for each of the regions with the new lists 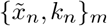. We label this new correlation function as the region-dependent component *ϕ*, reasoning that it captures rate correlation that can be attributed to the regional features, according to the model of Equation 5. We then extract *θ* from the total correlation using equation 6. The results of this decomposition are plotted in Figure 6b (a more detailed view in regions CGI, Enhancer, and SINE is shown in Supplemental Figure S2). We find that *ϕ* is more variable between different genomic regions, as compared to the component *θ*, which appears nearly uniform across genomic regions and in all regions shows rapid decay within ≈ 100 bp. We find that *θ* is generally well fit by a single-exponential decay and the fitted decay constants are similar across genomic regions, with a mean decay constant of 0.028 (ranging from .023 to .032, in TFBR and SINE, respectively), corresponding to a mean decay length of 36 basepairs (31 to 43 bp, respectively), Figure 6c. Importantly, this decomposition approach does not impose any *a priori* assumptions on the functional form of *θ*. The results confirm our hypothesis, that after removal of the correlation component dependent on regional features of the chromatin landscape, the remaining correlation decays exponentially with distance.

Our model predicts that the fitted exponential decay constant should be reflective of parameters of enzyme diffusion, namely, equal to 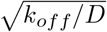. Thus, the model predicts that the decay constant is insensitive to other factors, such as variability of inter-CpG spacing. We confirm this in the experiment-derived correlation functions, by comparing the fitted decay lengths in each region to the inter-CpG distance distributions in each region (Figure 6d). Despite the significant differences in CpG density in the different regions, the decay lengths are generally constant, i.e., the median inter-CpG distance is 10 and 161 bp in CGI versus Intergenic regions, with corresponding fitted decay lengths of 39.6 and 35.6 bp, respectively.

Note that the exponential fit is not perfect, as evident by slight discrepancies in fit constants obtained for different window sizes. This discrepancy is due to low, persistent, nonzero longer-range correlation visible in *θ* in some regions (Figure S2), which we attribute to our model of *ϕ*, which was based only on three genomic features, and thus likely did not fully account for all region-based correlation. We therefore report fit constants from the 100bp window, to estimate the short-range decay while minimizing contamination from long-range residual correlation. However, the quantitative impact of different fit window sizes on estimated constants is relatively minor, as seen in Figure 6c.

All in all, these findings support that *θ* reflects the diffusion-based contribution to the correlation function, because (i) it is the explicitly distance-dependent part that remains after removing correlation attributed to the other features {*q*, *r*, *s*}_*m*_, (ii) it decays exponentially, (iii) it is uniform across the genome, consistent with processivity being an inherent property of DNMT1’s mode of action, and thus uniform across genomic regions. If *θ* is interpreted as reflecting processivity of the enzyme according to a 1D diffusion model, then diffusion parameters can be obtained from the experiment-derived correlation function. We thus estimate *D/k_off_* to be 1300 bp^2^ and the length of a processive step of DNMT1 to be 36 basepairs, on average, across the genome.

## DISCUSSION

### Summary of key results and methodological contribution

We have analyzed correlations among CpG sites in the genome obtained from estimated kinetics of post-replication DNA methylation and from WGBS in hESCs. We find that post-replication remethylation rates on nearby CpGs are correlated, and the nature of this correlation can shed light on molecular mechanisms of maintenance methylation, when analyzed in conjunction with stochastic simulation and mathematical models. We summarize our key results as follows: (i) Methylation rate correlation is a new genomic quantity, which contains information distinct from that contained in methylation state correlation. In particular, the rate correlation reveals mechanistic information on enzymatic processes. (ii) Some, but not all, of the rate correlation observed on nearby CpGs decays exponentially with genomic distance, and is consistent with processive activity of DNMT1 according to a linear diffusion model. Our analysis indicates an average distance of 36 bp between nearby CpGs that are methylated processively after DNA replication. (iii) In addition to evidence of processivity, we also discovered additional correlation, not consistent with a processive model, that is dependent on the genomic regional context. Among studied regions, CGIs and TFBRs showed the most significant contribution of this non-processive (termed “region-based”) correlation, which persisted past 1 kbp. In contrast, SINE and LINE showed the least contribution, suggesting that CpG interdependence in these regions results nearly entirely from linear diffusion (processivity) of the enzyme. (iv) Further analysis of the region-based rate correlation indicates that much of the regional variation can be attributed to variation in three genomic/epigenomic features: chromatin accessibility, CpG density, and bulk methylation.

More generally, we show how combining “top-down” mathematical modeling (i.e., data-driven, using statistical inference) with a “bottom-up” approach (i.e., using hypothesis-driven, or mechanistic models) can be used to glean kinetic insights from a measurement technique that affords genomewide readout of the post-replication methylome over time (32). A major new contribution of our paper is the development of a method for using data-derived rate correlations on genomic sites as a quantitative fingerprint of diffusive and nondiffusive enzyme kinetics. This method could be applied to other datasets in the future.

A number of experimental techniques have recently been developed, employing nucleoside-analog labeling of replicating DNA, followed by isolation/immunoprecipitation and sequencing (21, 38–47). Our study shows that statistical correlations from such measurements have potential to yield quantitative insights into molecular mechanisms governing epigenetic inheritance, even when time resolution is coarse.

Diffusion of proteins along DNA has been an area of intense study, notably in the context of how transcription factors locate target sites (reviewed in (37, 48)). Direct measurement of linear diffusion of proteins along DNA *in vivo* has been achieved by single-molecule tracking (49, 50). Our study shows that, for enzymes with ubiquitous target sites on DNA, correlations from temporal sequencing can also yield quantitative details on protein diffusion along DNA within cells (albeit indirectly), without recourse to labeling and microscopy.

### Processivity of DNMT1

While a number of studies have discovered evidence of DNMT1 processivity previously in *in vitro* systems (22–24, 28, 51, 52), our study is the first to our knowledge to uncover a quantifiable signature of methyl-transferase processivity in genome-wide, mammalian cell-based measurements. Thus our findings shed new light on enzymatic processivity within the context of maintenance methylation *in vivo*. Our findings are consistent with a picture wherein DNMT1 performs processive catalysis regardless of genomic context and in a quantitatively consistent manner (i.e., with relatively uniform lengths of processive steps between CpG targets, genome-wide). Our analysis does not afford direct estimation of linear coefficient *D*, but rather the ratio *D/k_off_*. We are not aware of any existing quantitative estimates or measurements of DNMT1 linear diffusion coefficient *in vivo*. If we assume that DNMT1 has *D* within similar range to other measured DNA-binding proteins (of order 10^5^ to 10^7^ bp ^2^/s (37, 53, 54)), our results would indicate average residence times of roughly 10^-4^ to 10^-1^ seconds for the enzyme when it is nonspecifically bound to non-CpG sites, en route to catalytic sites.

Previous *in vitro* estimates of the length of processive runs of DNMT1 varied widely. For example, Vilkaitis, et al reported processive runs as long as 520 bp (23) while Goyal et al. reported processive runs of over 6000 bp (28). Our estimates based on the Repli-BS dataset are much shorter than these, with the average length of a processive run being about 36 basepairs for nearby CpGs. One possible explanation for the discrepancy is that our estimate is from experiments in cells, where the *in vivo* chromatin environment, replication machinery, and full complement of DNA-binding proteins are present. These could limit free diffusion of DNMT1 along DNA. Transcription factors are also thought to search for targets partly by 1D diffusion along DNA, and the effect of crowding has been considered (37). While *in vitro* estimates for 1D sliding lengths of various DNA binding proteins are as high as 20 kb (53), *in vivo* sliding length of the lac repressor was found to be short, at 45 bp (49). How the chromatin environment affects the motion of eukaryotic DNA binding proteins is still poorly understood (55). In particular, the question of how methyltransferase processivity is affected by the crowded environment of the cell warrants further study.

Our analysis of the exponential contribution to rate correlation can not be directly or certainly attributed to the action of DNMT1 alone. For example, PCNA plays a role in recruitment of DNMT1 to replication foci(56), and this protein also can diffuse linearly along DNA (54) (although DNMT1 processivity does not rely on PCNA (23)). We cannot exclude the possibility that some mechanism other than intrinsic DNMT1 diffusion along DNA gives rise to the observed rate correlations, though this interpretation is consistent with the uniform lengthscale observed across different chromatin environments.

### Nonprocessive CpG coupling

We find that additional mechanisms affect post-replication remethylation rate correlation where the local genomic/chromatin landscape allows it. We find this non-processive (i.e. non-exponential) source of rate correlation to be most prevalent where chromatin is locally open, CpGs are dense, and the background methylation is relatively low. Although our study does not attempt to define the mechanistic basis of the non-processive component of the rate correlation, various mechanisms can be proposed based on our observations and on previous literature. For example, any mechanism whereby DNMT1 reaches its CpG target through cooperative interactions with other molecules could be speculated to have kinetics dependent on the local genomic and epigenomic context. Neighbor rate correlations could then be sensitive to local context and have lengthscales determined by the the cooperative molecular interactions, rather than being solely dependent on linear inter-CpG genomic distances, in contrast with the processive mechanism. For example, recruitment of DNMT1 to replicating DNA by UHRF1 (25, 26) likely results in context-dependent kinetics, since UHRF1 targeting is dependent on both histone state and the presence of hemi-methylated CpGs (57). A recent finding of monoubiquinated histone H3 helping recruit DNMT1 to DNA stretches with multiple, but not one, hemimethylated CpGs (20) supports the idea that UHRF1 helps direct DNMT1 to CpG-dense regions, and is consistent with our observation of higher rate correlation in CpG-dense regions. Our findings may also be consistent with a nucleation model, in which the initial binding of DNMT1 to replicating DNA occurs on nucleosomes, directed by UHRF1, after which DNMT1 reaches nearby CpGs processively (58). If the initial binding events of separate DNMT1s on nearby nucleosomes are correlated, such correlation would contribute on a lengthscale on the order of hundreds of basepairs, while shorter-lengthscale correlation would be introduced through processivity.

In addition to UHRF1-mediated mechanisms, additional factors are likely to play a role in the non-exponential correlation we observe. First, DNA is not one-dimensional; DNMT1 could reach nearby CpGs by facilitated diffusion (combining 1D diffusion along DNA with 3D diffusion in the nucleosol to nearby sites)(37), or by intersegmental transfer, similar to transcription factors (48). Our processive model assumes a 1D substrate, but our results hint at sensitivity of maintenance kinetics to 3D DNA structure in the weak appearance of peaks consistent with nucleosome spacing (Figure 1). Finally, transient binding of post-replication DNA by transcription factors could introduce correlation into maintenance methylation kinetics, as transcription factors could transiently block access to CpGs by DNMT1 and thus delay remethylation. Such a mechanism could explain why we observe the most pronounced region-based correlation in TF-BRs.

### Relationship between WGBS and rate correlation

Our study reveals a significant difference between the local correlation of methylation state versus rate. We interpret this result as being due to the different information content of the two: WGBS experiments capture reads that largely reflect the stable methylation landscape (though still partially influenced by replication-associated temporal variability (32)), while the Repli-BS-derived methylation rates reflect the transient dynamic processes occurring post-replication. Of note, there appears to be some relationship between the two, as we observed similar trends in region specificity in methylation state and rate (specifically in the region-specific, or non-processive, contribution to correlation). We furthermore note that, when we restrict the analysis to common sites (thus retaining CpGs with intermediate to high methylation levels, as rates are only inferrable for these sites), the similarity between rate- and state-correlation is increased (i.e., as in Figure 3 versus Figure 1, where, for example, the decay of bulk state-correlation in SINE is nearly as rapid as the decay of rate-correlation). This suggests partial cross-talk between the transient post-replication methylation events (including processivity) and the stable methylation landscape. Indeed, it was previously reported that CpG co-methylation decays within tens to hundreds of bp, with enzyme processivity proposed as its mechanistic origin (35).

However, our results also show that, in some regions (such as CGI), methylation state correlation is significantly longer-lived than the processive lengthscale and significantly higher than the rate correlation in total, and so the relationship between state- and rate-correlation is not directly clear. As our analysis is based only on replication-associated methylation reactions, it seems likely that our rate correlations are largely indicative of DNMT1-mediated maintence methylation, which cannot by itself shape the methylation landscape. The stochastic models of this paper focus only on sub-cellcycle timescales and methylation maintenance, and thus assume that the methylation landscape (i.e., state correlation) is a static property dictated by the parent-strand landscape at the time of DNA replication. However, previous methylation models suggest that differences in kinetic parameters, including maintenance kinetics as well as demethylating and *de novo* methylation, shape the global methylation landscape across mitotic cycles (19, 34). Our method for obtaining data-driven kinetic correlations could be useful in the future to further improve these types of mathematical models.

### Implications for stability of the methylation landscape

A number of mathematical modeling studies have provided support for the presence of interactions (also called “collaboration”) among CpGs in dynamic methylation processes, including in maintenance, de novo methylation, and demethylation reactions (19, 27, 30, 31). These models, in which a CpG is in some way affected by the state of nearby CpGs, built upon the earlier, so-called “standard” model, wherein each CpG was considered to be independently targeted by methyltransferases (59). Crucially, as the cited studies showed, these interactions provide the necessary nonlinearity to enable bistability in the dynamic system. That is, they enable the same family of “reader” and “writer” enzymes to simultaneously maintain distinct states of hyper- and hypo-methylation on groups of CpGs in different parts of the genome, thus mimicking observed methylation patterns. While these models tend to be phenomenological in nature (i.e., capturing dynamic phenomena without necessarily encoding detailed molecular mechanisms), processivity can be considered to be one type of molecular mechanism that contributes to inter-CpG interactions. Indeed, mathematical modeling also supports the idea that diffusive processivity enhances multi-generational stability of methylation patterns (28), just as does other mechanisms of CpG interaction (19). It follows that interdependence of CpG methylation kinetics, as quantified by correlation in this paper, has relevance to human ageing and disease, since instability of the methylation landscape has been linked to both(60, 61).

The mathematical model of Haerter et al. predicted that local (nearest-neighbor) interactions of methylating reactions was sufficient to achieve stable propagation of methylation states over multiple generations, though longer-range interactions were required for demethylating reactions (19). In the present study, we find that maintenance methylation occurs largely independently in regions that are CpG sparse and show low region-dependent correlation, such as LINE. A nearest-neighbor-only model is consistent with our findings in regions such as 5UTR and SINE, where the processive lengthscale is on the same order as the typical inter-CpG distance. In CGIs, the processive lengthscale is longer than the inter-CpG distance (median 10 bp in the analyzed data), suggesting that CpGs in CGIs effectively interact beyond nearest neighbors. Strong coupling of methylation in CGIs is consistent with faster maintenance kinetics in CpG-dense regions, as has recently been reported (21). TFBRs showed inter-CpG distances similar to the processive lengthscale, however here processivity is compensated by weak but longer range (region-based) coupling (also evident in CGIs). These findings may predict enhanced stability of methylation in TFBRs and CGIs across mitotic cycles, although better understanding of the interplay of correlation lengthscales for *de novo* and demethylating reactions is needed.

### Limitations of our study

Our study shows that novel experimental techniques that probe replication-associated dynamics genome-wide can yield surprisingly detailed dynamic insights, despite the limited time resolution. However, the statistical inference approach is nevertheless limited. First, each site-specific inferred rate constant is the result of a fit of the data to a stochastic Poisson process; undoubtedly this is a simplistic model for dynamics that could potentially be temporally complex in reality, e.g., non-exponential or even nonmonotonic. Thus, individual estimates can be error-prone for a number of reasons, from the inability of the simplistic model to capture complex dynamics, to the limited timeresolution or sampling depth for a given site, or all of the above. Because of the inherent difficulty in characterizing dynamics in detail at any individual CpG genome-wide, we focused here on general features of the rate correlation functions that appear robust across a given genomic region. In this way, the whole-genome nature of the data partially compensates for the sparse temporal resolution. We also ensured that our inference and analysis pipeline performed well on synthetic datasets, generated from simulations. Going forward, it may be possible to yield more detailed dynamic insights with deeper sampling and more fine-grained time resolution (as achieved in recent experiments (21)), which could enable further investigation into detailed, local kinetic correlation.

While we focus our models around the enzyme kinetics of DNMT1 (for simplicity and because it is the dominant methyltransferase responsible for carrying out maintenance activity), we acknowledge that our rate correlations are likely impacted by the presence of other (*de novo*) methyltransferases, which have been known to associate with highly methylated CpG-dense location (i.e., CGIs) (62). Although our mechanistic models rigorously incorporate 1D diffusion, they lack the dynamic interplay of *de novo*, maintenance, and demethylation reactions that has been studied in mathematical models previously (14, 19). Our approach could be applied to different cell lines in the future, e.g., with specific methyltransferases disrupted, to further disentangle the molecular basis of kinetic correlation.

## MATERIALS AND METHODS

### Methods overview

The methodology of this paper can be summarized as follows. We reanalyzed data from Whole Genome Bisulfite Sequencing (WGBS) (63) and Replication-association Bisulfite sequencing (Repli-BS) (64) in human Embryonic Stem Cells (hESCs) using a combination of data-driven statistical inference and hypothesis-driven stochastic modeling. First, Maximum Likelihood Estimation was used to infer per-CpG post-replication remethylation rates from Repli-BS experiments, following our previously developed method (33). We analyzed the correlation of these data-inferred rates on nearby CpGs in different genomic contexts, such as Enhancer, Promoter, etc., to study regional differences in maintenance kinetics. Next, we studied the association of the strength of nearby-CpG remethylation-rate correlations with other local genomic/epigenomic features. To aid interpretation of the experiment-derived correlation functions and their regional differences and associations, we developed stochastic models of post-replication DNA methylation maintenance kinetics. Using these models, we generated simulated bisulfite sequencing datasets under different mechanistic hypotheses, and we compared the resultant *in silico*-generated rate correlation functions to experimental data.

### Site-specific post-replication methylation kinetics inference

The post-replication methylation data (Repli-BS data) of Human Embryonic Stem Cells (HUES 64) were downloaded from GSE82045. In the Repli-BS experiment, cells were pulsed for one hour with bromodeoxyuridine (BrdU). Then, bisulfite sequencing of BrdU-labeled DNA captured CpG methylation reads from DNA that was replicated during the pulse interval. The pulse-chase experiment captured methylation level of CpGs at timepoints 0, 1, 4, and 16 hours post-pulse, thereby giving a genome-wide temporal readout of CpG-methylation over sixteen hours postreplication.

The Maximum Likelihood Estimation (MLE) procedure for inferring per-CpG post-replication methylation rates from Repli-BS data is described in detail in (33). Briefly, the temporally distributed binary read-data (methylated-1 or unmethylated-0) at each CpG site is fitted by a Poisson process, with each site *i* characterized by two inferred constants, *k_i_* and *f_i_*, which represent the rate at which methylation accumulates at the site over the course of the experiment, and the steady-state (or long-time) fraction of cells in the measurement set that exhibit methylation at site *i*, respectively. We hereon refer to the inferred parameters *k_i_* as the “postreplication remethylation rates” or simply the “remethylation rates”. Details of the inference approach can be found in the Supplement (Extended Methods). Note that, while the persite inferences are obtained based on an analytical, independent Poisson process model, the inferred rate parameters can nevertheless be used to investigate more complex types of dynamics and inter-site dependencies through correlations that are observed among inferred parameters on nearby CpGs.

The ability to infer a remethylation rate for a given CpG site, and the uncertainty associated with that inferred rate, depends on the read-depth of the experimental data, which varies across sites and across timepoints. Details of uncertainty quantification can be found in the Supplement and in our previous study (33). We estimated on average 30% error in any given estimate of rates *k_i_*. Despite these relatively high error-rates, we validated our method by ensuring that ground-truth rate correlations (obtained from simulated data, see below), could be accurately recovered by the MLE inference pipeline.

### Annotations of genomic regions

The GRCh37/hg19 genome was used as the reference genome in this paper. The region annotations for genes, promoters, exons, introns, 3’UTRs and 5’UTRs are downloaded from the UCSC Genes track in UCSC Table Browser, whereas the LINEs(Long Interspersed Nuclear Elements), SINEs(Short Interspersed Nuclear Elements), LTRs(Long Terminal Repeats) were extracted from RepeatMasker track. CGIs(CpG Islands), Enhancers were downloaded from CpG Island track and Gene-Hancer track, respectively. The promoters in this paper were defined as regions 2000bp upstream and 200bp downstream of transcription start sites (TSS). Local CpG density for a site was defined as the number of neighboring CpGs in a 500bp window centered at that site.

The chromatin accessibility data was retrieved from [ENCODE/OpenChrom](http://hgdownload.cse.ucsc.edu/goldenPath/hg19/encodeDCC/wgEncodeOpenChromDnase/)(Duke University) H1 cell line. The histone modification peaks were downloaded from [ENCODE/Broad](http://hgdownload.cse.ucsc.edu/goldenPath/hg19/encodeDCC/wgEncodeBroadHistone/). The regions of transcription factor(TF) peaks or TFBR denoted in this paper were acquired from [ENCODE ChIP-seq clusters](https://hgdownload.cse.ucsc.edu/goldenpath/hg19/encodeDCC/wgEncodeRegTfbsClustered/) for 161 TFs in H1 cells.

### Stochastic simulation

Region-specific simulations of single-CpG stochastic enzyme-kinetic models were carried out using two candidate mechanisms, the Distributive model and the Processive model (Figure 4). The model reactions and associated rate parameters are graphically depicted in Figures 4b and 4c. The Distributive model was simulated using the Gillespie Stochastic Simulation Algorithm (65). To incorporate 1D diffusion into the Processive model, we used a First Passage Time Kinetic Monte Carlo algorithm inspired by (66). The methylation maintenance model and simulation method is based in part on our previous simulation studies (33). In the present paper, we refined our Processive model and simulation algorithm to rigorously incorporate physics of 1D diffusion (also called “sliding”) of proteins along DNA, with unbinding (37), while enabling simulation of large numbers of CpGs. Our analytical results on enzyme diffusion and simulation algorithm are presented in the Supplement (Extended Methods), along with further details of the models. Parameter values are chosen to be in line with experimentally measured values for DNMT1 where possible (67), and also to match features of the Repli-BS data (see Supplemental Table S1 for details). Simulation codes in MATLAB are available in a github repository: Read-Lab-UCI/DNA-methylation-kinetics-correlation.

Simulations are performed for stretches of *N* CpGs, (*N* = 25000 – 75000). Simulations mimic two types of experimental data modalities: WGBS and Repli-BS. To simulate WGBS experiments, for a region of CpGs, the simulation is initialized at post-replicaton time=0, and then read out at randomly sampled timepoints between zero and 24 hours later, to reflect the variable post-replication timings of bulk cells in WGBS experiments. Ten simulation replicates are combined to generate estimates of average per-site methylation levels. To mimic the Repli-BS experiment, the methylation status of CpGs in the simulation were read out at timepoints sampled from intervals matching pulse-chase experiments (64), including uncertainty with respect to true postreplication timing. That is, given the finite BrdU pulse-length, the 0-hour experimental timepoint is assumed to correspond to *t* ∈ [0 – 1] hours post-replication, the 1-hour experimental timepoint corresponds to *t* ∈ [1 – 2] hours postreplication, etc. Therefore, timepoints of simulation readout were sampled from *t_chase_* + *r* ∈ [0, 1] hour, where experimental timepoints *t_chase_* were 0,1,4,16 hours and *r* is a uniformly distributed random number. The number of simulations and read-outs at each timepoint were chosen to mimic the distribution of experimental read-depths. The synthetic Repli-BS data were then processed with the same MLE procedure as the experimental data, to infer per-CpG methylation rates. Simulations were used to validate our statistical inference procedure. We tested that ground-truth correlation functions produced by the two models could be recapitulated by the inference procedure.

## ACKNOWLEDGEMENTS

We thank Jun Allard for comments on the manuscript. This work was supported by NSF 2022182 to TD and ER, NSF DMS1715455 to ER, NSF DMS1763272, and a grant from the Simons Foundation (594598 QN).

## Supplemental Materials

### Extended Methods

#### A. Maximum Likelihood Estimation of Remethylation Rates

The Repli-BS dataset(32) contains read-depths that vary widely both across CpGs and across measured timepoints. The data can be expressed as 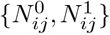, or the number of unmethylated (0) and methylated (1) reads observed at site *i* at timepoint *j*. We assume that the probability of finding a methylated read at site *i* and at time *t_j_* is

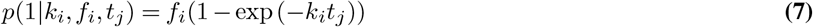

where *k_i_* is the rate of accumulation of methylation post-replication and *f_i_* is the steady-state fraction of cells in the population methylated at site *i*. Parameters are estimated from the data by Maximum Likelihood Estimation (MLE), as in (33). In the MLE pipeline, we use a per-site minimum read-depth threshold to increase the baseline accuracy of inferred parameters. In the current work, the threshold was set as follows: each CpG required a minimum of 5 reads (methylated or unmethylated) at both timepoint *t* = 0 and combined timepoints *t* > 0 (for a total minimum of 10 reads over all timepoints). Furthermore, each CpG required a minimum of 5 methylated reads in total (over all measured timepoints). (We performed detailed testing and uncertainty quantification for various methods of setting thresholds previously (33), finding that a read-depth-based threshold gave good balance between genomic coverage and uncertainty in estimates). For each site, we compute the Likelihood Function, *L*(*θ*||*x_i_*), where *θ* contains the parameters *θ* = {*k*, *f*} and *x_i_* is the observed data at site *i*, given the Poisson process model. We observed three cases: (1) the parameters are identifiable. This occurs when the likelihood function contains a global maximum in *θ* = {*k*, *f*}; the location of the maximum gives the inferred parameter values. The majority of sites in the dataset fall in this category; (2) The parameters are practically unidentifiable. This occurs when a global maximum exists, but the confidence interval is very wide. We discard sites for which this occurs, requiring that the 75% confidence interval on *k* (obtained by the profile likelihood method) fall within the range of values [0.01 – 10]h^-1^; (3) The rate *k* is unidentifiable, but its value can be bounded. This occurs frequently, as many sites show full methylation already at the first experimental timepoint, indicating a post-replication methylation rate that is faster than the time-resolution of the experiment, given the one-hour pulse length. We observe this in a likelihood function *L*(*θ*||*x_i_*) that plateaus at high rate values. In these cases, we use a procedure to estimate a lower bound for *k_i_*: we determine the 25% lower bound of the confidence interval on *k*, relative to the plateaued value of *L*(*θ*||*x_j_*). We choose this value for the lower bound on *k*, reasoning that it estimates the location where the likelihood function is no longer increasing significantly.

#### B. Stochastic Simulation

The models are written in the form of stochastic biochemical kinetic reaction models (or in the case of the processive mechanism, a stochastic reaction-diffusion model). The models concentrate solely on the process of maintenance methylation, i.e., the remethylation process that occurs in less than 24 hours, while ignoring other processes such as demethylation. Both models treat DNA as a 1D strand with CpG sites that can be unmethylated (u), hemi-methylated (h), or methylated (m). Sites are assumed to be unmethylated or hemimethylated immediately after replication, with hemimethylated sites susceptible to remethylation by the enzyme. The distributive and processive models differ in how the enzyme moves to new hemimethylated sites after catalyzing methylation at one site. The Distributive mechanism treats individual CpG sites as fully independent. The DNMT1 will be disassociated from DNA after it methylated a hemimethylated site. By contrast, in the Processive model, after methylation DNMT1 can diffuse towards its neighbor sites either upstream or downstream. We assume the enzyme travels with 1D diffusion coefficient D and unbinds with rate koff. *E* denotes a free enzyme (not bound to DNA). *Eh* is a complex formed when *E* is bound to *h* (similar for *m*). The model reactions are as follows.

##### B.1. Distributive Model

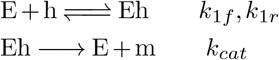

After the catalytic step, unbinding is assumed to happen rapidly, therefore the catalytic step and unbinding are assumed to occur together in one step, with rate *k_cat_* (i.e., we assume that *k_off_* ≪ *k_cat_*).

##### B.2. Processive Model

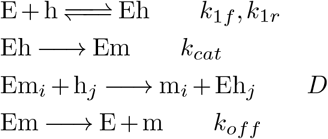

In the processive model, the diffusive step allows the enzyme to reach a hemimethylated CpG from a nearby CpG in *Em* state. The likelihood of this occuring before unbinding with rate *k_off_* depends on the properties of diffusion (see below).

##### B.3. Model Parameters

Model parameters were chosen to be approximately in line with experimentally measured values for DNMT1 where possible (67), while also showing reasonable agreement with our experiment-derived methylation rates and rate correlation (in the case of the Processive model). Parameters used in simulations are shown in Table 1.

##### B.4. Simulation algorithms

For the distributive model, the standard Gillespie Stochastic Simulation Algorithm was used (65). For the processive model, we combine stochastic reactions with a 1D lattice model of diffusion. The lattice spaciging is 1 basepair. We apply a First Passage Time Kinetic Monte Carlo algorithm inspired by (66). This allows us to simulate diffusive steps efficiently, since we do not need to track each diffusive hop with basepair resolution. Rather, the algorithm requires sampling from the First Passage Time Distribution for an enzyme to reach a nearby CpG by diffusion. This allows us to simulate large stretches (tens of thousands) of CpGs.

In the simulations, the enzyme is allowed to diffuse bidirectionally along DNA. Consider an enzyme with a starting position located at a CpG in the Em state. Let there be two nearest neighbor CpGs in the *h* state (which are thus candidates to be subsequently targeted by the enzyme, according to the Processive model reactions above). We refer to the diffusion *Domain* as the 1D region between left and right neighbors. Let *d_L_* be the distance (in bp) between the left neighbor and the starting position, and similarly *d_R_* for the right neighbor. Then the next event that can occur for this Em CpG is one of: (1) the enzyme diffuses distance *d_L_* to the left neighbor with diffusion coefficient D. The current CpG undergoes Em → m and the left neighbor undergoes h → Eh; (2) Similar for the right neighbor; or (3) The enzyme unbinds from DNA before reaching either left or right neighbor at rate *k_off_*, so Em → E+m. For the simulation, we must compute the waiting time distribution (or First Passage Time Distribution *f*(*t*)) for the enzyme to exit the domain by one of these events. We must also compute the exit probabilities, i.e. the probability of events (1),(2) or (3) occuring, which we refer to as *P_L_*, *P_R_*, *P_off_*, and *P_L_* + *P_R_* + *P_off_* = 1. The distribution *F*(*t*) and the exit probabilities can be solved analytically (see Finite Domain model, below). These events, which involve diffusion, have non-exponential waiting times, and we refer to them as *FPT* (first passage time) events. The other reactions (*k*_1*f*_, *k*_1*r*_, *k_cat_*) are assumed to proceed via standard Markovian (memoryless) kinetics, with exponential waiting times, and are referred to as *KMC* (kinetic Monte Carlo) events. Following (66), a sketch of the algorithm for the processive simulation is then:

Initialize *N* CpGs at post-replication time *t_p.r_*. =0 in *u* or *h* state by sampling from input methylation landscape

**while** *t_p.r._* < *t_out_* (*t_out_* is the desired readout time)

*Determine which CpG undergoes next event and which reaction occurs at that site by:*

* For all CpGs in h or Eh, determine next *KMC* event and time to event, *τ_KMC_*, by the Gillespie algorithm (65).
* *Build new diffusion domains*. For all CpGs newly in Em state, determine domain length *L*=*d_R_* + *d_L_* and *x*_0_ = *d_L_*. Sample 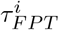 from *F*(*t*) and store in event queue. Compute corresponding exit probabilities for each *i*th site, 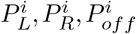.
* *Update diffusion domains*. For all CpGs remaining in Em state since last event, determine whether left or right neighbor has changed. If so, rebuild diffusion domain and resample 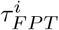 from *F*(*t*) and replace in event queue.
* Select the minimum *FPT* time, and the corresponding site *j* for the next *FPT* event, 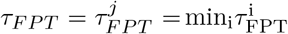.
**if** *τ_KMC_* < *τ_FPT_*, set *τ_next_* = *τ_KMC_*. Update system state according to *KMC* event.
**else** *τ_next_* = *τ_FPT_*. Sample from the exit probabilities for the *j*th site to determine outcome and update system state according to *FPT* event.
**end if**
Update the system time: *t_p.r._* = *t_p.r._* +*τ_next_*.
Update *FPT* queue by 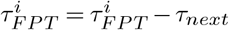. (Remove the site which underwent diffusion previously from event queue).
**end while**

Record system state.

**Table S1.**
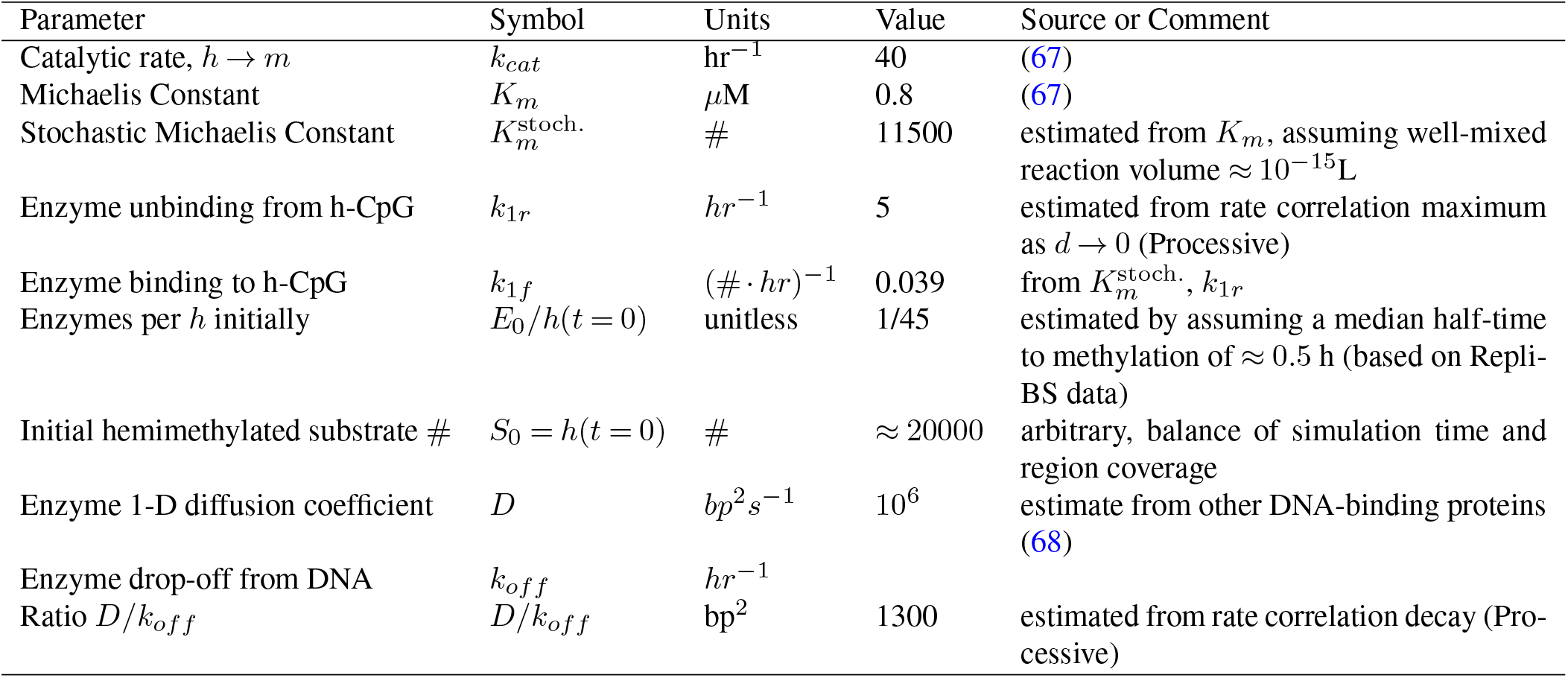
Table of Parameters

#### C. Analytic 1D Diffusion Model

##### C.1. Semi-infinite Domain

Following the model described in the main text, consider two CpG sites, linear distance *d* apart. Let *P*(*x*, *t*) be the probability density of a particle (enzyme) to be at location *x* in 1D at time *t*. For simple diffusion (no unbinding) the governing equation

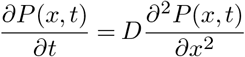

with initial condition *P*(*x* = 0, *t* = 0) = 1 and absorbing boundary condition *P*(*x* = *d*,*t*> 0) = 0 gives the density

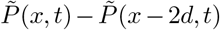

where 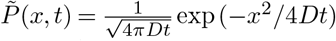 is the solution to the standard 1D diffusion model on an infinite domain (no absorbing boundary condition).

When unbinding is included in the model, the governing equation becomes:

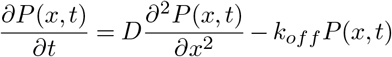

The solution to this PDE with the same absorbing condition at distance *d* as above gives

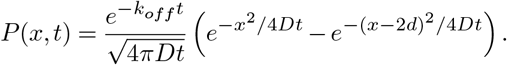

We then have the survival probability for the protein to remain bound on DNA at time *t*:

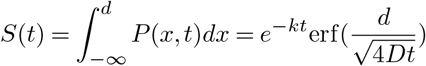

leading to an expression for the total probability flux to “exit” the DNA (either by unbinding or by absorption at *x* = *d*):

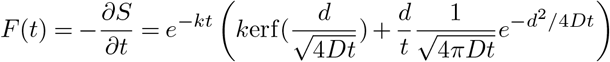

The first term is the flux to exit by unbinding, the second is the flux to the absorbing state at distance *d*, i.e. *F*(*t*) = *F_unbind_*(*t*) + *F_absorb_*(*t*). Total probability to reach the absorbing state at distance *d* by diffusion, integrating the absorbing flux,

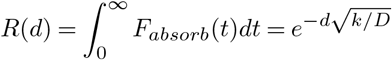

Thus, the probability for the enzyme to reach a site at distance *d* away from the initial site before unbinding is given by 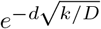.

##### C.2. Finite Domain

In the simulations, we need to consider not just pairs of CpGs distance *d* apart, but consider that an enzyme bound to DNA may have two neighbors within reach in *h* state, left and right. Let the total domain length be *L*=*d_R_* + *d_L_*, and the enzyme starting position is *x*_0_ = *d_L_*. We have a diffusion model, with unbinding, with initial condition *P*(*x* = *x*_0_,*t* = 0) = 1 and absorbing boundary conditions at both ends, *P*(*x* = 0,*t*> 0) = 0; *P*(*x* = *L*,*t*> 0) = 0. The Fourier series solution is

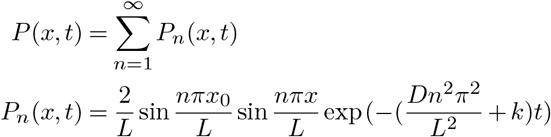

The survival probability of the enzyme on DNA at location *x* (i.e. before either unbinding or reaching left or right neighbor) is

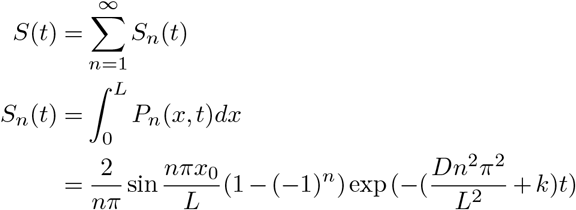

The first passage time distribution *f*(*t*) can be obtained by normalizing the time-dependent probability flux 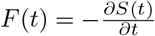. For purposes of the simulation, we sample from *f*(*t*) making use of the C.D.F. for *f*(*t*). This C.D.F. is given by:

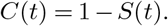

To determine the exit probabilities as a function of time, (i.e., the relative likelihood of exiting left, right, or to solution), we make use of the time-dependent probability fluxes to each end-state: *F_L_*(*t*), *F_R_*(*t*), *F_solution_*(*t*). These are obtained as:

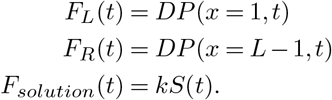

#### D. Relationship between Semi-infinite domain model and Processive simulation

Using the analytical semi-infinite domain model together with the arguments in main text, the correlation between methylation times at two sites separated by distance *d* is given by 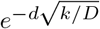. In our Processive simulation, there are a number of additional complicating factors that are not present in the idealized theoretical model. For example, after the enzyme reaches the neighbor by diffusion, there is a chance for it to unbind (rate *k*_1*r*_) before successfully catalyzing methylation (rate *k_cat_*). There is also a chance that the neighbor site will already be occupied by another copy of the enzyme. Finally, in the simulation *τ_D_* is on the same order as *τ_i_*, not much less as assumed in the idealized model, because it accounts for the fast diffusion time as well as the slower time for complex formation and methylation, introducing delay between subsequent methylation events. All of these factors serve to decrease the correlation at a given distance *d*, but can be assumed to not significantly affect the distance-dependence, thus predicting that correlation should be proportional, but not strictly equal to, 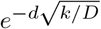.

Note the distinction between the correlation of methylation times at individual sites (on which our analytical theory is based is based) and methylation rates, which we extract from experiments and simulations. Exact methylation times at individual sites are not currently measureable in genome-wide pulse-chase experiments, but methylation rates *k_i_* are estimated from Repli-BS-Seq experiments, where, for a given CpG sites *i*, the experiment-inferred methylation rate constant *k_i_* is by definition taken to be equal to 1/ < *τ_i_* >, i.e., the inverse of the average over methylation times at site *i* across different cells. The inverse relationship between rates and times could in principle affect the linear Pearson correlation, however, from simulations we found that the Pearson correlation function was not sensitive to the distinction, and the rate correlation decay was well described by 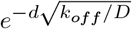, in accordance with the analytical theory (see Figure S1).

## Supplementary Figures

**Fig. S1.**
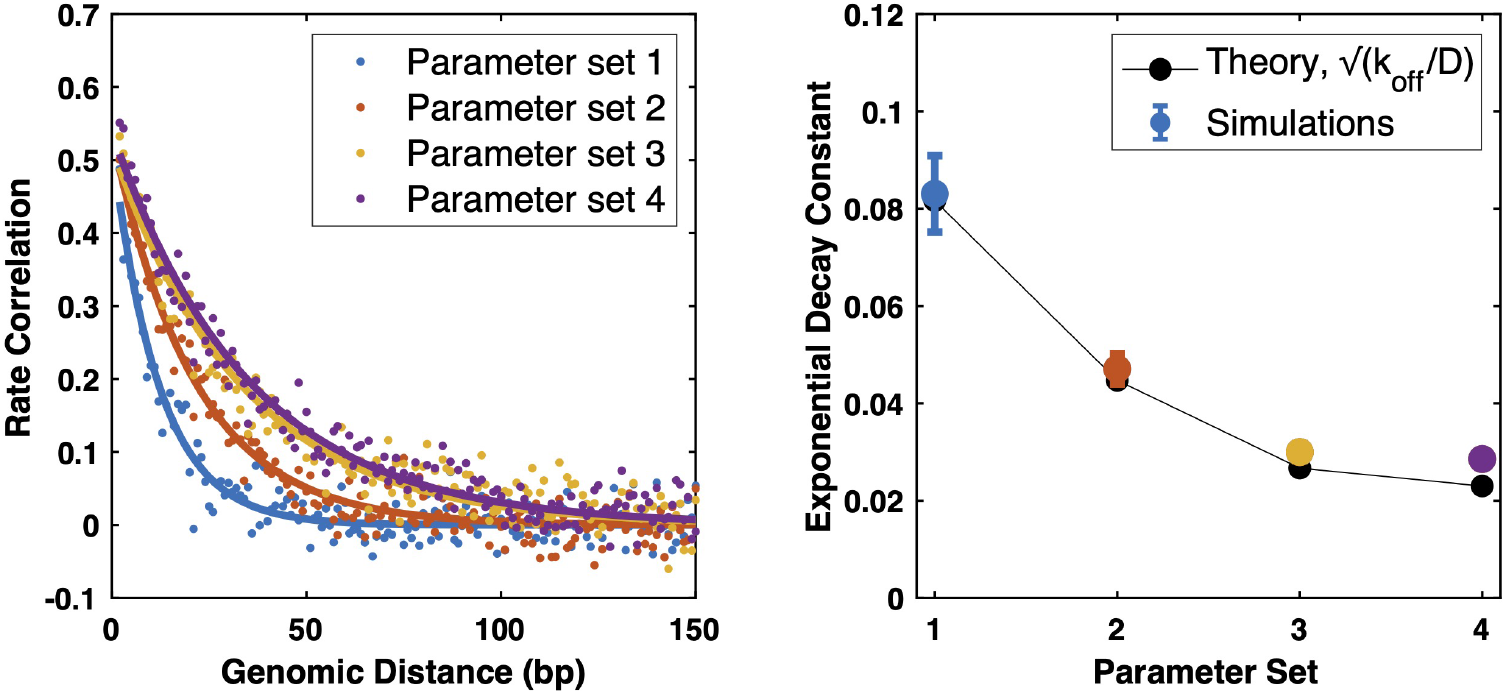
Comparison of analytical theory of enzyme processivity to stochastic simulations. (Left) Processive simulations were performed choosing four different values for the parameter ratio D/koff. (Values for sets {1,2,3,4} were {150,500,1402,1890}*bp*^2^. All other parameters were held constant.) After inferring remethylation rates from the synthetic data by the MLE pipeline, ensuing rate correlations were fitted to a single exponential decay function in an unbiased manner. (Right) Fitted decay constants from simulated data (colored dots) overlaying the theoretically predicted decay constant from the model parameters 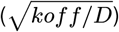, showing close agreement of theory and simulation, despite the increased complexity of the methylation dynamics simulation, as compared to the 1D analytical model.

**Fig. S2.**
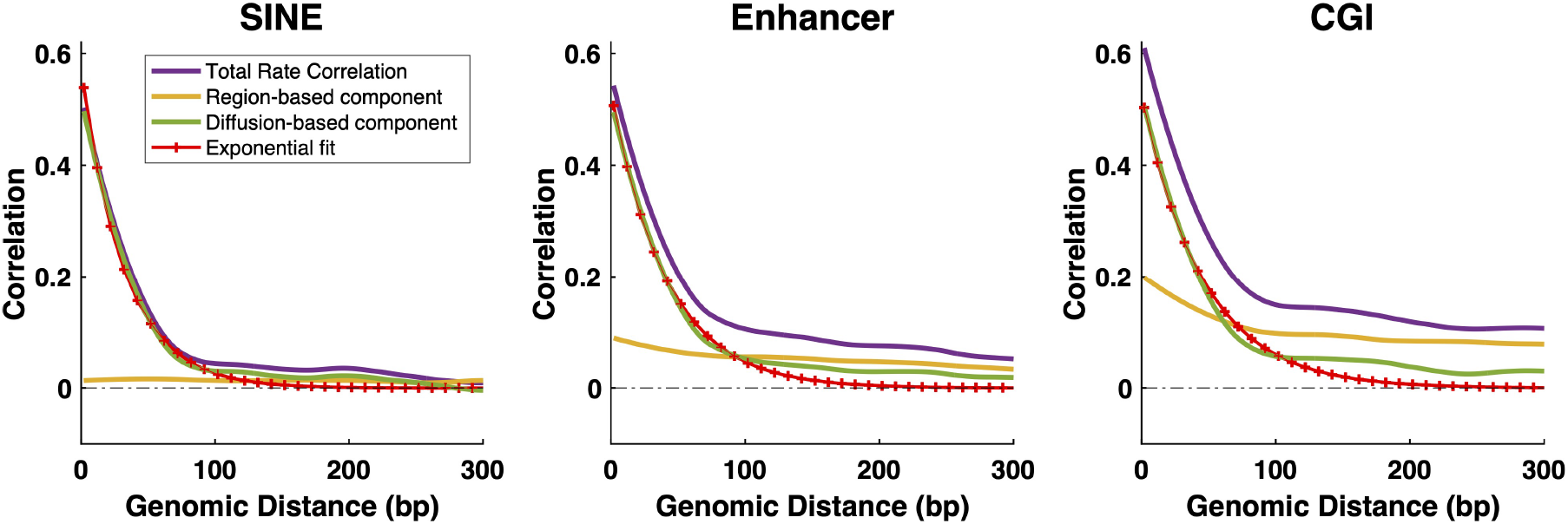
Decomposition of experiment-derived rate correlation functions by region into region-based and diffusion-based components. Simultaneous plotting of correlation components and exponential fits for the three representative regions (same data as Figure 6b in main text, but providing a more detailed view). SINE shows a nearly-zero region-based component (yellow), indicating that the total rate-correlation is already reasonably well-fit by an exponential decay. In contrast, Enhancer and CGI both show small but persistent region-based correlation that is the dominant contributor to the total correlation beyond 100 bp (Enhancer) and beyond 50 bp (CGI). Removal of the region-based component according to the model in Figure 6 results in remaining component *θ* (green), which is reasonably well fit by a single exponential decay. However, some residual nonzero correlation is still apparent at longer range in the green curves. The exponential fits shown correspond to a 100 bp fitting window to fit the short-range decay while minimizing contribution from residual correlation at longer range.

## Reference

1. Suhua Feng, Steven E Jacobsen, and Wolf Reik. Epigenetic reprogramming in plant and animal development. Science, 330(6004):622–627, 2010.

2. Zachary D Smith and Alexander Meissner. Dna methylation: roles in mammalian development. Nature Reviews Genetics, 14(3):204–220, 2013.

3. Adrian Bird. Dna methylation patterns and epigenetic memory. Genes & development, 16 (1):6–21, 2002.

4. Albert Jeltsch. Beyond watson and crick: Dna methylation and molecular enzymology of dna methyltransferases. Chembiochem, 3(4):274, 2002.

5. Robert J Klose and Adrian P Bird. Genomic dna methylation: the mark and its mediators. Trends in biochemical sciences, 31(2):89–97, 2006.

6. Michela Curradi, Annalisa Izzo, Gianfranco Badaracco, and Nicoletta Landsberger. Molecular mechanisms of gene silencing mediated by dna methylation. Molecular and cellular biology, 22(9):3157–3173, 2002.

7. Vishal Nanavaty, Elizabeth W Abrash, Changjin Hong, Sunho Park, Emily E Fink, Zhuangyue Li, Thomas J Sweet, Jeffrey M Bhasin, Srinidhi Singuri, Byron H Lee, et al. Dna methylation regulates alternative polyadenylation via ctcf and the cohesin complex. Molecular cell, 78(4):752–764, 2020.

8. Peter A Jones and Shirley M Taylor. Cellular differentiation, cytidine analogs and dna methylation. Cell, 20(1):85–93, 1980.

9. Arthur D Riggs. X inactivation, differentiation, and dna methylation. Cytogenetic and Genome Research, 14(1):9–25, 1975.

10. Peter A Jones and Gangning Liang. Rethinking how dna methylation patterns are maintained. Nature Reviews Genetics, 10(11):805–811, 2009.

11. En Li, Timothy H Bestor, and Rudolf Jaenisch. Targeted mutation of the dna methyltransferase gene results in embryonic lethality. Cell, 69(6):915–926, 1992.

12. Robin Holliday and John E Pugh. Dna modification mechanisms and gene activity during development. Science, 187(4173):226–232, 1975.

13. Timothy H Bestor. The dna methyltransferases of mammals. Human molecular genetics, 9 (16):2395–2402, 2000.

14. Albert Jeltsch and Renata Z Jurkowska. New concepts in dna methylation. Trends in biochemical sciences, 39(7):310–318, 2014.

15. Matthew C Lorincz, Dirk Schübeler, Shauna R Hutchinson, David R Dickerson, and Mark Groudine. Dna methylation density influences the stability of an epigenetic imprint and dnmt3a/b-independent de novo methylation. Molecular and cellular biology, 22(21):7572–7580, 2002.

16. Yingying Zhang, Christian Rohde, Sascha Tierling, Tomasz P Jurkowski, Christoph Bock, Diana Santacruz, Sergey Ragozin, Richard Reinhardt, Marco Groth, Jörn Walter, et al. Dna methylation analysis of chromosome 21 gene promoters at single base pair and single allele resolution. PLoS genetics, 5(3):e1000438, 2009.

17. Gilad Landan, Netta Mendelson Cohen, Zohar Mukamel, Amir Bar, Alina Molchadsky, Ran Brosh, Shirley Horn-Saban, Daniela Amann Zalcenstein, Naomi Goldfinger, Adi Zundelevich, et al. Epigenetic polymorphism and the stochastic formation of differentially methylated regions in normal and cancerous tissues. Nature genetics, 44(11):1207–1214, 2012.

18. Julia Arand, David Spieler, Tommy Karius, Miguel R Branco, Daniela Meilinger, Alexander Meissner, Thomas Jenuwein, Guoliang Xu, Heinrich Leonhardt, Verena Wolf, et al. In vivo control of cpg and non-cpg dna methylation by dna methyltransferases. PLoS genetics, 8 (6):e1002750, 2012.

19. Jan O Haerter, Cecilia Lövkvist, Ian B Dodd, and Kim Sneppen. Collaboration between cpg sites is needed for stable somatic inheritance of dna methylation states. Nucleic acids research, 42(4):2235–2244, 2014.

20. Yuichi Mishima, Laura Brueckner, Saori Takahashi, Toru Kawakami, Junji Otani, Akira Shinohara, Kohei Takeshita, Ronald Garingalao Garvilles, Mikio Watanabe, Norio Sakai, et al. Enhanced processivity of dnmt1 by monoubiquitinated histone h3. Genes to cells, 25(1): 22–32, 2020.

21. Xuan Ming, Zhuqiang Zhang, Zhuoning Zou, Cong Lv, Qiang Dong, Qixiang He, Yangyang Yi, Yingfeng Li, Hailin Wang, and Bing Zhu. Kinetics and mechanisms of mitotic inheritance of dna methylation and their roles in aging-associated methylome deterioration. Cell Research, 30(11):980–996, 2020.

22. Andrea Hermann, Rachna Goyal, and Albert Jeltsch. The dnmt1 dna-(cytosine-c5)-methyltransferase methylates dna processively with high preference for hemimethylated target sites. Journal of Biological Chemistry, 279(46):48350–48359, 2004.

23. Giedrius Vilkaitis, Isao Suetake, Saulius Klimašauskas, and Shoji Tajima. Processive methylation of hemimethylated cpg sites by mouse dnmt1 dna methyltransferase. Journal of Biological Chemistry, 280(1):64–72, 2005.

24. Željko M Svedružić and Norbert O Reich. Mechanism of allosteric regulation of dnmt1’s processivity. Biochemistry, 44(45):14977–14988, 2005.

25. Magnolia Bostick, Jong Kyong Kim, Pierre-Olivier Estève, Amander Clark, Sriharsa Pradhan, and Steven E Jacobsen. Uhrf1 plays a role in maintaining dna methylation in mammalian cells. Science, 317(5845):1760–1764, 2007.

26. Jafar Sharif, Masahiro Muto, Shin-ichiro Takebayashi, Isao Suetake, Akihiro Iwamatsu, Takaho A Endo, Jun Shinga, Yoko Mizutani-Koseki, Tetsuro Toyoda, Kunihiro Okamura, et al. The sra protein np95 mediates epigenetic inheritance by recruiting dnmt1 to methylated dna. Nature, 450(7171):908–912, 2007.

27. Laura B Sontag, Matthew C Lorincz, and E Georg Luebeck. Dynamics, stability and inheritance of somatic dna methylation imprints. Journal of theoretical biology, 242(4):890–899, 2006.

28. Rachna Goyal, Richard Reinhardt, and Albert Jeltsch. Accuracy of dna methylation pattern preservation by the dnmt1 methyltransferase. Nucleic acids research, 34(4):1182–1188, 2006.

29. Shijie C Zheng, Martin Widschwendter, and Andrew E Teschendorff. Epigenetic drift, epigenetic clocks and cancer risk. Epigenomics, 8(5):705–719, 2016.

30. You Song, Honglei Ren, and Jinzhi Lei. Collaborations between cpg sites in dna methylation. International Journal of Modern Physics B, 31(20):1750243, 2017.

31. Loukas Zagkos, Mark Mc Auley, Jason Roberts, and Nikos I Kavallaris. Mathematical models of dna methylation dynamics: Implications for health and ageing. Journal of theoretical biology, 462:184–193, 2019.

32. Jocelyn Charlton, Timothy L Downing, Zachary D Smith, Hongcang Gu, Kendell Clement, Ramona Pop, Veronika Akopian, Sven Klages, David P Santos, Alexander M Tsankov, et al. Global delay in nascent strand dna methylation. Nature structural & molecular biology, 25 (4):327–332, 2018.

33. Luis Busto-Moner, Julien Morival, Honglei Ren, Arjang Fahim, Zachary Reitz, Timothy L Downing, and Elizabeth L Read. Stochastic modeling reveals kinetic heterogeneity in postreplication dna methylation. PLOS Computational Biology, 16(4):e1007195, 2020.

34. Paul Adrian Ginno, Dimos Gaidatzis, Angelika Feldmann, Leslie Hoerner, Dilek Imanci, Lukas Burger, Frederic Zilbermann, Antoine HFM Peters, Frank Edenhofer, Sébastien A Smallwood, et al. A genome-scale map of dna methylation turnover identifies site-specific dependencies of dnmt and tet activity. Nature communications, 11 (1):1–16, 2020.

35. Shicheng Guo, Dinh Diep, Nongluk Plongthongkum, Ho-Lim Fung, Kang Zhang, and Kun Zhang. Identification of methylation haplotype blocks aids in deconvolution of heterogeneous tissue samples and tumor tissue-of-origin mapping from plasma dna. Nature genetics, 49(4):635–642, 2017.

36. William G Jacoby. Loess:: a nonparametric, graphical tool for depicting relationships between variables. Electoral Studies, 19(4):577–613, 2000.

37. Leonid Mirny, Michael Slutsky, Zeba Wunderlich, Anahita Tafvizi, Jason Leith, and Andrej Kosmrlj. How a protein searches for its site on dna: the mechanism of facilitated diffusion. Journal of Physics A: Mathematical and Theoretical, 42(43):434013, 2009.

38. R Scott Hansen, Sean Thomas, Richard Sandstrom, Theresa K Canfield, Robert E Thurman, Molly Weaver, Michael O Dorschner, Stanley M Gartler, and John A Stamatoyannopoulos. Sequencing newly replicated dna reveals widespread plasticity in human replication timing. Proceedings of the National Academy of Sciences, 107(1):139–144, 2010.

39. Tyrone Ryba, Dana Battaglia, Benjamin D Pope, Ichiro Hiratani, and David M Gilbert. Genome-scale analysis of replication timing: from bench to bioinformatics. Nature protocols, 6(6):870–895, 2011.

40. Claire Marchal, Takayo Sasaki, Daniel Vera, Korey Wilson, Jiao Sima, Juan Carlos Rivera-Mulia, Claudia Trevilla-García, Coralin Nogues, Ebtesam Nafie, and David M Gilbert. Genome-wide analysis of replication timing by next-generation sequencing with e/l repliseq. Nature protocols, 13(5):819–839, 2018.

41. Qian Du, Saul A Bert, Nicola J Armstrong, C Elizabeth Caldon, Jenny Z Song, Shalima S Nair, Cathryn M Gould, Phuc-Loi Luu, Timothy Peters, Amanda Khoury, et al. Replication timing and epigenome remodelling are associated with the nature of chromosomal rearrangements in cancer. Nature communications, 10(1):1–15, 2019.

42. Chenhuan Xu and Victor G Corces. Genome-wide mapping of protein-dna interactions on nascent chromatin. Methods in molecular biology (Clifton, NJ), 1766:231, 2018.

43. Constance Alabert, Teresa K Barth, Nazaret Reverón-Gómez, Simone Sidoli, Andreas Schmidt, Ole N Jensen, Axel Imhof, and Anja Groth. Two distinct modes for propagation of histone ptms across the cell cycle. Genes & development, 29(6):585–590, 2015.

44. Nazaret Reverón-Gómez, Cristina González-Aguilera, Kathleen R Stewart-Morgan, Nataliya Petryk, Valentin Flury, Simona Graziano, Jens Vilstrup Johansen, Janus Schou Jakobsen, Constance Alabert, and Anja Groth. Accurate recycling of parental histones reproduces the histone modification landscape during dna replication. Molecular cell, 72(2):239–249, 2018.

45. Pauline Vasseur, Saphia Tonazzini, Rahima Ziane, Alain Camasses, Oliver J Rando, and Marta Radman-Livaja. Dynamics of nucleosome positioning maturation following genomic replication. Cell reports, 16(10):2651–2665, 2016.

46. Mónica P Gutiérrez, Heather K MacAlpine, and David M MacAlpine. Nascent chromatin occupancy profiling reveals locus-and factor-specific chromatin maturation dynamics behind the dna replication fork. Genome research, 29(7):1123–1133, 2019.

47. Chenhuan Xu and Victor G Corces. Nascent dna methylome mapping reveals inheritance of hemimethylation at ctcf/cohesin sites. Science, 359(6380):1166–1170, 2018.

48. Jason Gorman and Eric C Greene. Visualizing one-dimensional diffusion of proteins along dna. Nature structural & molecular biology, 15(8):768–774, 2008.

49. Petter Hammar, Prune Leroy, Anel Mahmutovic, Erik G Marklund, Otto G Berg, and Johan Elf. The lac repressor displays facilitated diffusion in living cells. Science, 336(6088):1595–1598, 2012.

50. Kiyoto Kamagata, Yuji Itoh, Cheng Tan, Eriko Mano, Yining Wu, Sridhar Mandali, Shoji Takada, and Reid C Johnson. Testing mechanisms of dna sliding by architectural dnabinding proteins: dynamics of single wild-type and mutant protein molecules in vitro and in vivo. Nucleic acids research, 49(15):8642–8664, 2021.

51. Timothy H Bestor and Vernon M Ingram. Two dna methyltransferases from murine erythroleukemia cells: purification, sequence specificity, and mode of interaction with dna. Proceedings of the National Academy of Sciences, 80(18):5559–5563, 1983.

52. Sabrina Adam, Hiwot Anteneh, Maximilian Hornisch, Vincent Wagner, Jiuwei Lu, Nicole E Radde, Pavel Bashtrykov, Jikui Song, and Albert Jeltsch. Dna sequence-dependent activity and base flipping mechanisms of dnmt1 regulate genome-wide dna methylation. Nature communications, 11(1):1–15, 2020.

53. Kiyoto Kamagata, Eriko Mano, Kana Ouchi, Saori Kanbayashi, and Reid C Johnson. High free-energy barrier of 1d diffusion along dna by architectural dna-binding proteins. Journal of molecular biology, 430(5):655–667, 2018.

54. Anna B Kochaniak, Satoshi Habuchi, Joseph J Loparo, Debbie J Chang, Karlene A Cimprich, Johannes C Walter, and Antoine M van Oijen. Proliferating cell nuclear antigen uses two distinct modes to move along dna. Journal of Biological Chemistry, 284(26):17700–17710, 2009.

55. David M Suter. Transcription factors and dna play hide and seek. Trends in cell biology, 30 (6):491–500, 2020.

56. Linda S-H Chuang, Hang-In Ian, Tong-Wey Koh, Huck-Hui Ng, Guoliang Xu, and Benjamin FL Li. Human dna-(cytosine-5) methyltransferase-pcna complex as a target for p21waf1. Science, 277(5334):1996–2000, 1997.

57. Xiaoli Liu, Qinqin Gao, Pishun Li, Qian Zhao, Jiqin Zhang, Jiwen Li, Haruhiko Koseki, and Jiemin Wong. Uhrf1 targets dnmt1 for dna methylation through cooperative binding of hemimethylated dna and methylated h3k9. Nature communications, 4(1):1–13, 2013.

58. Robert M Vaughan, Bradley M Dickson, Matthew F Whelihan, Andrea L Johnstone, Evan M Cornett, Marcus A Cheek, Christine A Ausherman, Martis W Cowles, Zu-Wen Sun, and Scott B Rothbart. Chromatin structure and its chemical modifications regulate the ubiquitin ligase substrate selectivity of uhrf1. Proceedings of the National Academy of Sciences, 115 (35):8775–8780, 2018.

59. Sarah P Otto and Virginia Walbot. Dna methylation in eukaryotes: kinetics of demethylation and de novo methylation during the life cycle. Genetics, 124(2):429–437, 1990.

60. Adiv A Johnson, Kemal Akman, Stuart RG Calimport, Daniel Wuttke, Alexandra Stolzing, and Joao Pedro De Magalhaes. The role of dna methylation in aging, rejuvenation, and age-related disease. Rejuvenation research, 15(5):483–494, 2012.

61. Atsuya Nishiyama and Makoto Nakanishi. Navigating the dna methylation landscape of cancer. Trends in Genetics, 2021.

62. Jing Liao, Rahul Karnik, Hongcang Gu, Michael J Ziller, Kendell Clement, Alexander M Tsankov, Veronika Akopian, Casey A Gifford, Julie Donaghey, Christina Galonska, et al. Targeted disruption of dnmt1, dnmt3a and dnmt3b in human embryonic stem cells. Nature genetics, 47(5):469–478, 2015.

63. Bradley E Bernstein, John A Stamatoyannopoulos, Joseph F Costello, Bing Ren, Aleksandar Milosavljevic, Alexander Meissner, Manolis Kellis, Marco A Marra, Arthur L Beaudet, Joseph R Ecker, et al. The nih roadmap epigenomics mapping consortium. Nature biotechnology, 28(10):1045–1048, 2010.

64. Jocelyn Charlton, Timothy Downing, Zachary Smith, Hongcang Gu, Kendell Clement, Ramona Pop, Veronika Akopian, Sven Klages, David Santos, Alexander Tsankov, Bernd Timmermann, Michael Ziller, Evangelos Kiskinis, Andreas Gnirke, and Alexander Meissner. Global delay in nascent strand dna methylation. Nature Structural & Molecular Biology, 25 (4):327–332, 2018.

65. Daniel T Gillespie. Exact stochastic simulation of coupled chemical reactions. The journal of physical chemistry, 81(25):2340–2361, 1977.

66. Andri Bezzola, Benjamin B Bales, Richard C Alkire, and Linda R Petzold. An exact and efficient first passage time algorithm for reaction–diffusion processes on a 2d-lattice. Journal of Computational Physics, 256:183–197, 2014.

67. Ivan Hemeon, Jemy A Gutierrez, Meng-Chiao Ho, and Vern L Schramm. Characterizing dna methyltransferases with an ultrasensitive luciferase-linked continuous assay. Analytical chemistry, 83(12):4996–5004, 2011.

68. Anahita Tafvizi, Fang Huang, Alan R Fersht, Leonid A Mirny, and Antoine M van Oijen. A single-molecule characterization of p53 search on dna. Proceedings of the National Academy of Sciences, 108(2):563–568, 2011.

